# Comparing cellular response to two radiation treatments based on key features visualization

**DOI:** 10.1101/2024.02.29.582706

**Authors:** Polina Arsenteva, Olivier Guipaud, Vincent Paget, Morgane Dos Santos, Georges Tarlet, Fabien Milliat, Hervé Cardot, Mohamed Amine Benadjaoud

## Abstract

**Motivation:** In modern treatment by radiotherapy, different irradiation modalities can be used, potentially producing different amounts of adverse effects. The differences between these modalities are often studied via two-sample time course in vitro experiments. The resulting data may be of high complexity, in which case simple methods are unadapted for extracting all the relevant information.

**Methods:** In this article we introduce network-based tools for the visualization of the key statistical features, extracted from the data. For the key features extraction we utilize a statistical framework performing estimation, clustering with alignment of temporal omic fold changes originating from two-sample time course data.

**Results:** The approach was applied to real transcriptomic data obtained with two different types of irradiation. The results were analyzed using biological literature and enrichment analysis, thus validating the robustness of the proposed tools as well as achieving better understanding of the differences in the impact of the treatments in question.

**Availability and implementation:** Python package freely available here: https://github.com/parsenteva/scanofc.

## 1 Introduction

Radiotherapy is a one of the main types of cancer treatment, along with surgery and chemotherapy, received by approximately 60% of all cancer patients (Warren et al., 2008). It is based on using ionizing radiation to either kill cancer cells or block their ability to divide. Similarly to other types of treatment, radiotherapy may induce adverse effects, that is undesirable changes to healthy tissues situated around the irradiated tumor. Modern technological advances give us access to a vast range of different modes of irradiation, that may vary in terms of dose, volume, energy, etc. It is of high interest for potential clinical applications to be able to choose such modes of radiotherapy that minimize the amount of potential adverse effects.

In this paper, we study the difference between two modes of radiotherapy, differing in energy levels, through comparing the in vitro gene expressions of human endothelial cells. Within the vascular network, the endothelium, made up of a monolayer of endothelial cells at the interface between the blood and the tissues, plays a major role in the response of normal tissues and tumors to radiotherapy (Guipaud et al., 2018; Wijerathne et al., 2021). Damage to endothelial cells by ionizing radiation is associated with late intestinal and lung dysfunctions (Rannou et al., 2015; Toullec et al., 2017), justifying the interest in better understanding the continuum and dynamics of molecular changes that propagate over time and lead to tissue damage. Endothelial cells could therefore represent one of the main cellular constituents of organs at the origin of the initiation and development of late radiation-induced lesions, which makes it an interesting target for the prediction, prevention and management of radiation damage. Efforts have been made to understand the molecular response of these cells at different molecular scales by multi-omics approaches, generating a wealth of valuable data to go further in the use of endothelial cells as a predictive tool (Heinonen et al., 2015; Jaillet et al., 2017; Heinonen et al., 2018; Benadjaoud et al., 2022; Morilla et al., 2022). In this sense, we have established proofs of concept that multiparametric measurements on these cells make it possible to highlight differences in response between irradiation modalities differing by the energy or the dose rate of the radiation (Paget et al., 2019; Ben Kacem et al., 2020, 2022).

In order to study the dynamic of the response to irradiation under different conditions, we measure the gene expression for multiple time points. These data are rich in information but also complex to analyze from the statistical point of view. Indeed, the data in questions has four dimensions: number of genes, number of time points, the former being significantly larger than the latter, two experimental conditions, and the number of replicates. The last dimension is necessary in order to account for the uncertainty, due to the destructive nature of measurements, which makes it impossible to observe an actual temporal signal based on one replicate only.

The most widely used tool in radiobiology to compare two types of treatment is relative biological effectiveness, or RBE (Valentin, 2003). Currently, RBE is derived from the linear quadratic model which relates the absorbed dose to the fraction of surviving cells by performing a clonogenic assay (Munshi et al., 2005). This approach provides important information but remains too simplistic to accurately predict radio-induced adverse effects. The goal of this paper is to propose multi-parametric alternatives to RBE that are capable of representing the complex data in question, as means to quantitatively and qualitatively compare different irradiation modalities.

In this work, we apply the mathematical framework proposed by Arsenteva et al. (2023), with the goal of extracting key features from each of the considered datasets corresponding to a distinct irradiation modality. The framework of interest consists in modeling the radio-induced temporal fold changes for all the considered genes, and performing their clustering jointly with temporal alignment, in the way that takes into account the correlations between genes, the measurement uncertainties, as well as the destructive nature of the measurements. Based on this formalism, and inspired by gene regulatory networks (Riccadonna et al., 2016; Nguyen and Braun, 2018), we propose two tools for the visualization of the objects encoding the key features of the considered datasets: microscopic and mesoscopic fold change networks. We perform the enrichment analysis of the obtained gene clusters, which allows to validate our approach to key feature visualization as a predictive tool.

## 2 Materials and methods

### 2.1 Experimental protocol

#### 2.1.1 Establishment of a time-course transcriptomic signature of irradiation

We designed a custom TaqMan Low-Density Array (TLDA) (Thermo Fisher Scientific) to measure the expression levels of 192 mRNAs involved in the response of HUVECs to ionizing radiation (Table S1). For the most part, the 192 genes were chosen based on our previous work. We have also identified some genes of interest based on knowledge in the field of radiobiology and by exploiting the results of the enrichments we carried out on the proteomic and transcriptomic datasets. The way this transcriptomic signature of ionizing radiation was established is described below. First, we considered data from proteomic analyses performed on proteins from irradiated HUVECs (Morilla et al., 2022). Data was obtained by iTRAQ-2D-LC-MS/MS after 20 Gy irradiation, which allowed quantification and identification of 2000-3000 proteins per time post-irradiation at 8 time points (12 hours, 1, 2, 3, 4, 7, 14 and 21 days). For each time point, we considered proteins for which there is a statistically significant difference between irradiated and non-irradiated samples. We then selected the proteins for which the fold change is greater than or equal to +/- 1.5 compared to the non-irradiated samples, and then we determined the number of times each of these candidates is differentially expressed in different time intervals after irradiation. Finally, we selected proteins that are differentially expressed at 3 or more time points in the 0.5-7 day time interval (65 proteins) and at 2 or more time points in the 7-21 day time interval (68 proteins), representing a total of 59 unique proteins (Table S2). Secondly, we considered the data from targeted transcriptomic analyzes obtained on HUVECs irradiated at a single dose of 2 or 20 Gy, or at a fractionated dose of 20 Gy (10 × 2 Gy per day for 10 days). Data were generated by RT-qPCR using a panel of predefined TLDAs targeting angiogenesis, inflammation, apoptosis, immune response and protein kinase functions (Heinonen et al., 2015, 2018; Morilla et al., 2022), and a premade RT^2^ Profiler™ PCR array targeting protein glycosylation functions (Jaillet et al., 2017). All these data allowed the quantification of 517 mRNAs per time point over 8 time points (12 hours, 1, 2, 3, 4, 7, 14 and 21 days). In the same way as for the selection of candidate genes from the proteomic study, we selected the genes for which the fold change is statistically significantly greater than or equal to +/- 2 compared to unirradiated samples at each time post-irradiation (adjusted p-value < 0.05). We then determined the number of times each of these candidates is differentially expressed in different time intervals after irradiation, and we selected the genes that are differentially expressed at 2 or more time points in the time interval 7 to 21 days (68 genes), and at 3 or more time points in the time intervals 0.5 to 7 days (65 genes) and 0.5 to 21 days (67 genes), representing a total of 94 unique genes. Similarly, 13 genes involved in protein glycosylation were selected based on the results of the RT^2^ Profiler™ PCR array. A total of 107 unique genes were finally selected from the transcriptomic analysis dataset (Table S3). Third, we performed sub-network enrichment analyses of common regulators of the 164 unique entities selected from transcriptomic and proteomic datasets (59 proteins and 107 genes, 2 common entities between the 2 sets) using the Pathway Studio Web Mammal software, version 12.4, querying the Mammal (Anatomy; CellEffect; DiseaseFx; GeneticVariant; Viruses) version 12.4.0.3 (Updated April 25, 2021) database from Elsevier (www.elsevier.com/pathway-studio) (Nikitin et al., 2003). This allowed the identification and selection of 11 additional genes potentially involved in the response to ionizing radiation. At last, we removed a few genes from the proteomic analysis to reach the number of 184 genes on the custom TLDA, and we added a few genes of interest based on our published work on radiation-induced endothelial-to-mesenchymal transition (EndoMT) (Mintet et al., 2015, 2017) (12 genes), endothelial senescence (Soysouvanh et al., 2020; Benadjaoud et al., 2022) (3 genes) and chemokines involved in the response to radiation (5 gens) (Tables S4-S9). Overall, after considering common genes between different datasets, this signature includes 184 genes. We added 5 reference genes, one of which (18S rRNA) is replicated 4 times on the TLDA plate (required by the manufacturer) (Table S1 and Figure S1).

#### 2.1.2 Endothelial cell culture, irradiation procedure, RNA isolation and reverse transcription and quantitative real time PCR

Human umbilical vascular endothelial cells (HUVECs) from Lonza were cultured in EGM-2-MV medium at 37°C with 5% CO2. To compare the effects of the different ionizing radiation beam energy spectra on the transcriptomic response of HUVECs over time, we measured transcriptional profiles with real time qPCR under control and under a single irradiation dose of 20 Gy at 0 h using either a LINAC (at 4 MV, 2.5 Gy/min) or a SARRP (at 220 kV, 2.5 Gy/min) with measurements at 2, 4, 7, 14 and 21 days. Four biological replicate cell populations were separated from a single population just prior to experiments. Non-irradiated cells (control samples) were left for the same amount of time on a benchtop near the irradiation room. All irradiations were carried out at 90–100% cell confluence and the same number of population doublings (passage 3, corresponding to 9 to 12 population doublings). For long term experiments (7–21 days post-irradiation), culture medium was changed every week. Total RNAs were prepared using the mirVana miRNA isolation kit (Thermo Fisher Scientific). RNA concentrations were quantified on a NanoDrop ND-1000 apparatus (NanoDrop Technologies). Reverse transcription was performed with 2 µg RNA using the High-Capacity Reverse Transcription Kit (Thermo Fisher Scientific). cDNA (700 ng) per sample was loaded onto the port of each gene signature array cards and PCR was performed with the ABI PRISM 7900 Sequence detection system (Applied Biosystems). RQ Manager and Data Assist software (Thermo Fisher Scientific) were used to determine the 2^−ΔΔCt^values for each of the genes of each replicate with fixed criteria: a maximum allowable Ct value at 37 was fixed and maximum Ct values were not included in calculations. Normalization was performed using a global normalization method (Mestdagh et al., 2009), i.e. the software first finds the common assays among all samples and the median Ct of those assays is used as the normalizer, on a per sample basis. Hereafter, the datasets combining the generated 2-ddCT values are named LINAC dataset and SARRP dataset depending on the radiation source used.

### 2.2 Key features extraction

We apply the framework presented in Arsenteva et al. (2023) in order to obtain gene clusters for datasets LINAC and SARRP. Using the same notations, we denote *n*_*e*_ the number of genes observed for a given dataset, *p* the number of time points where the measurements are taken, and *n*_*r*_ the number of replicates for a given time point. Clustering is performed with respect to the temporal fold changes of genes. The fold changes encode the difference in response between the irradiated and non-irradiated experimental conditions for a given gene over time, and are assumed to be Gaussian, i.e. for the gene 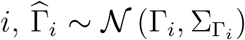. The mean is the pointwise estimator 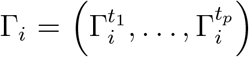, such that 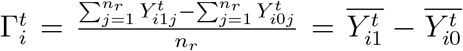, with 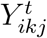 the response of gene *i* at time point *t* under the experimental condition *k* (1 if irradiated and 0 otherwise) and replicate *j*. The covariance 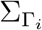 is a diagonal matrix, with the diagonal 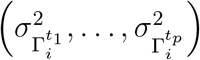, such that 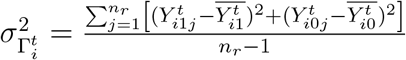

For the purpose of aligning the fold changes in the multivariate temporal setting, Arsenteva et al. (2023) introduce a transformation 𝒲_*s*_, which is a sort of a multivariate time warp of step *s* ∈ ℤ, transforming a pair of fold changes 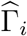 and 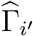 into their translated versions 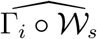 and 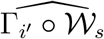. A dissimilarity measure used to compare aligned fold changes is defined as follows:

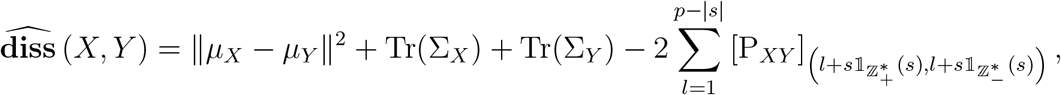

given Gaussian vectors *X* and *Y* with respective means µ_*X*_ and µ_*Y*_, variance matrices Σ_*X*_ and Σ_*Y*_, and cross-covariance matrix P_*XY*_. This dissimilarity measure allows to perform the comparison based on means as well as measurement uncertainties and correlations between genes contained in the covariance matrix, on the fold changes having undergone a translation of step *s*.

The algorithm introduced in Arsenteva et al. (2023) performs k-medoids clustering based on the Optimal Warping Dissimilarity matrix, defined as

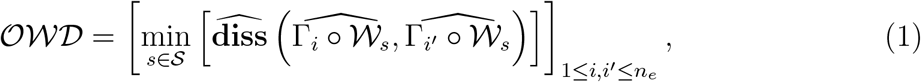

making it a computationally efficient equivalent of performing the fold changes clustering iteratively with alignment. For a given fixed number of clusters *K*, the algorithms returns the following quantities:

- clusters, denoted *cluster*_*k*_ = {*i* ∈ {1, …, *n*_*e*_}|*Cl*_*i*_ = *k*} for *k* ∈ {1, …, *K*}, where 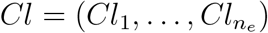 are the cluster labels for all genes,
- centroids *C* = (*C*_1_, …, *C*_*K*_), i.e. the most representative genes for each cluster,
- warps for all entities 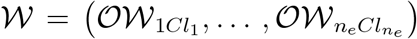, representing the number of steps necessary to reach the optimal alignment with the corresponding centroid, calculated based on the Optimal Warp matrix defined as follows:

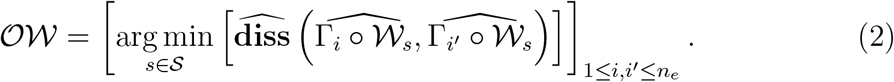

Arsenteva et al. (2023) propose an additional feature of sign-penalizing the dissimilarity between the fold changes. This is particularly pertinent in the context treated in this paper, since it is of interest to distinguish genes that are up-regulated as a result of radiation treatment from those that are down-regulated.

### 2.3 Network inference for key features visualization

#### 2.3.1 Microscopic network

A classical version of the network of omic fold changes (also referred to as microscopic to distinguish from the mesoscopic network representation discribed in the next section) is modeled by a random graph 𝒢 = (𝒩, ℰ), where 𝒩 is the node set, and ℰ is the edge set. The set 𝒩 consists of genes and is of size *n*_*e*_, whereas the set ℰ represents connections between the genes. The graph is described by the binary adjacency matrix 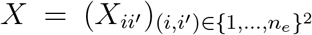 such that *X*_*ii*′_ = 1 denotes the existence of an edge between entities *i* and *i*′, and 0 denotes the absence of one.

The adjacency matrix for omic fold changes is constructed from a similarity measure that is formulated based on the dissimilarity between fold changes estimators. In its simplest form it is calculated based on the Optimal Warping Dissimilarity matrix:

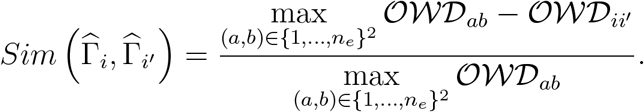

It can also be reformulated in a minimax form, which makes apparent its link with the dissimilarity:

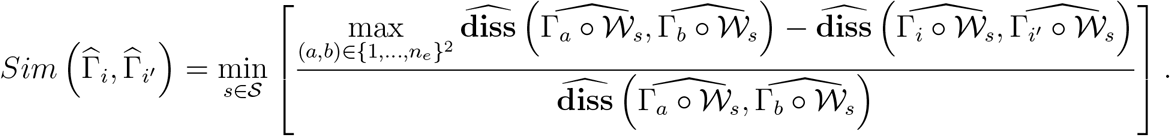

It can be noted that the formulation of the similarity measure in the case without time warping is achieved through the same definition by considering the degenerate warping space 𝒮 = {0}. In all cases, this similarity measure is bounded by 0 from below and 1 from above, the value in case of *i* = *i*′ being 1 and in case of 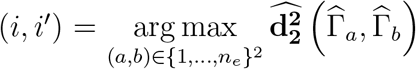 being 0, which makes the measure more easily interpretable and comparable for different omic datasets.

Let us denote a set of all unique entity pairs as *pairs* = {(*i, i*′) ∈ {1, …, *n*_*e*_}^2^|*i* < *i*′}. We define the empirical cumulative distribution function 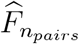 of similarity over the observed fold change pairs as follows:

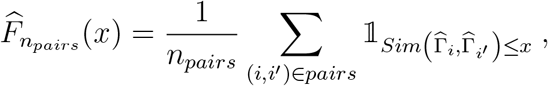

where 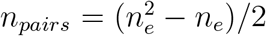, and *x* ∈ [0, 1] is a similarity level. In other words, 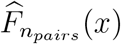 represents the proportion of fold change pairs less or as similar as *x*. For **p** ∈ [0, 1], representing sparsity level of the network, an empirical **p**-quantile is constructed as follows:

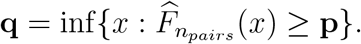

In order to define the elements of the adjacency matrix 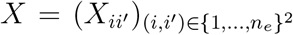, we distinguish two cases. If 𝒢 is undirected, its elements are equal to

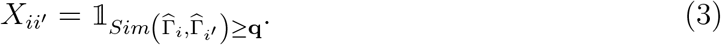

This definition implies that all entities that are at least as similar as the quantile corresponding to the chosen network sparsity level will be considered as connected, and not connected otherwise. In case if 𝒢 is directed, the elements are defined as

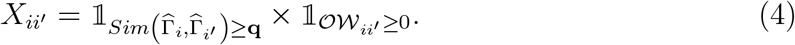

The additional term means that only predictive and simultaneous relations with respect to pairwise warps from 𝒪𝒲 remain in the directed case. Matrix 𝒪𝒲 can be naturally used in this context since it is anti-symmetric, as stated in Proposition 2.2 of Arsenteva et al. (2023). The adjacency matrix is thus no longer symmetric: the symmetry is preserved for non-significant (absent) connections and simultaneous (bidirectional) connections, the remaining connections being anti-symmetric.

#### 2.3.2 Mesoscopic network

This section introduces a type of network representation that combines key features obtained with both fold changes clustering and network inference described above. The mesoscopic network is modeled by a random graph 𝒢_ℳ_ = (𝒩_ℳ_, ℰ_ℳ_), where 𝒩_ℳ_ is the node set, and ℰ_ℳ_ is the edge set. As in the microscopic case, the node set 𝒩_ℳ_ contains biological entities, but in this case it is of size *K*, corresponding to the number of clusters, labeled by centroids *C* = (*C*_1_, …, *C*_*K*_). The remaining information on clustering is contained in the associated node set weight function, defined as follows:

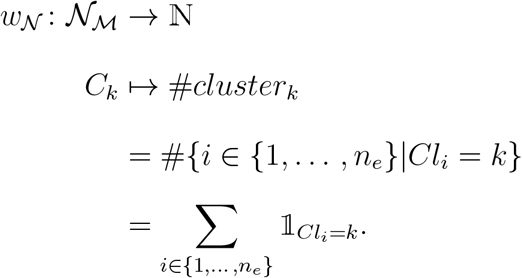

The edge set contains the main information on connections between the clusters in the form of distributions, encoded in the associated edge set weight function. The definition of the weight function differs in the undirected and the directed cases. The undirected case is defined below:

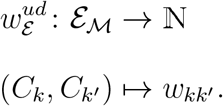

where *w*_*kk*′_ defines edge thickness and encodes the total number of connections between the given pair of clusters:

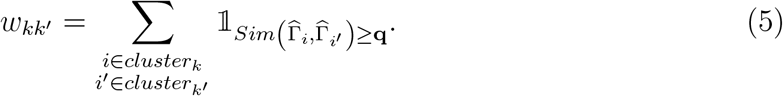

The directed case requires an extended version of the weight function, adding arrowheads on both sides of the edge:

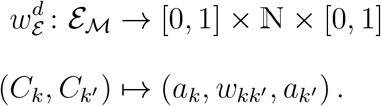

where *a*_*k*_ and *a*_*k*′_ define the sizes of the arrowheads towards *C*_*k*_ and *C*_*k*′_ encoding the proportions of the predictive connections in the corresponding directions. The formal definitions are given below:

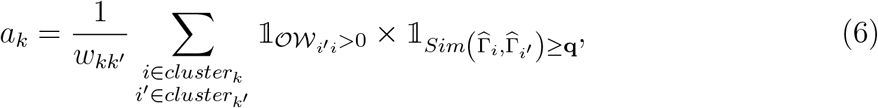

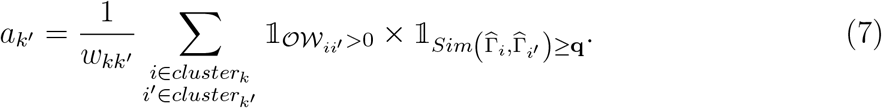

It can be noticed that the quantities given in (5), (6) and (7) can be expressed only based on the elements of the adjacency matrix, defined in (4), instead of both the similarity and the 𝒪𝒲 matrix. In particular, the expression for the edge thickness can be expressed as

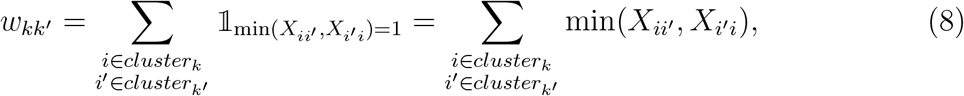

using the fact that if, for a given pair of entities *i* ∈ *cluster*_*k*_ and *i*′ ∈ *cluster*_*k*′_ for *k* ≠ *k*′, we have *Sim* 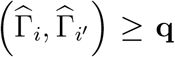, then either *X*_*ii*′_ or *X*_*i*′*i*_ or both are equal to 1. We apply a similar reasoning to rewrite the expression for the arrowhead sizes, utilizing the symmetry of *Sim*(·, ·) and the anti-symmetry of 𝒪𝒲:

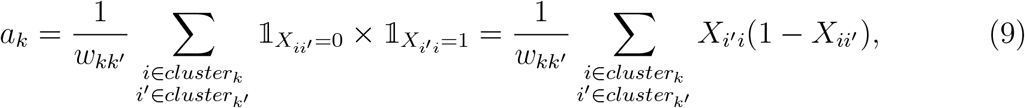

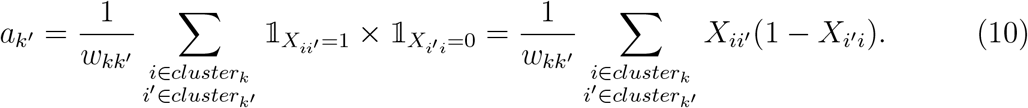

Indeed, considering the equations (6) and (9), saying that *Sim* 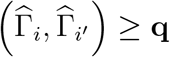 and 𝒪𝒲_*i*′*i*_ > 0 is equivalent to saying that *Sim* 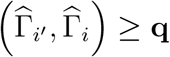 and 𝒪𝒲_*ii*′_< 0, which is true if and only if *X*_*ii*′_ = 0 and *X*_*i*′*i*_ = 1, based on (4).

#### 2.3.3 Illustrative examples

In the package *ScanOFC*, we propose visualizations for the microscopic and mesoscopic fold changes networks. Figures 1 and 2 illustrate examples of such networks, both calculated based on the adjacency matrix, presented in Supplementary Table S10. In this toy example there are 11 entities (e.g. genes), labeled with numbers from 1 to 11. We suppose that the entities are distributed in 3 clusters: *cluster*_1_ = {1, 2, 3}, *cluster*_2_ = {4, 5, 6} and *cluster*_3_ = {7, 8, 9, 10, 11}, with the corresponding centroids *C*_1_ = 1, *C*_2_ = 4 and *C*_3_ = 7. It can be noted that the microscopic network representation in Figure 1 has a block structure, with blocks corresponding to clusters, and with the centroid nodes bigger than the others. The visualization also distinguishes between the two types of connections: green edges correspond to predictive connections, whereas gray to simultaneous ones.

**Figure (1).**
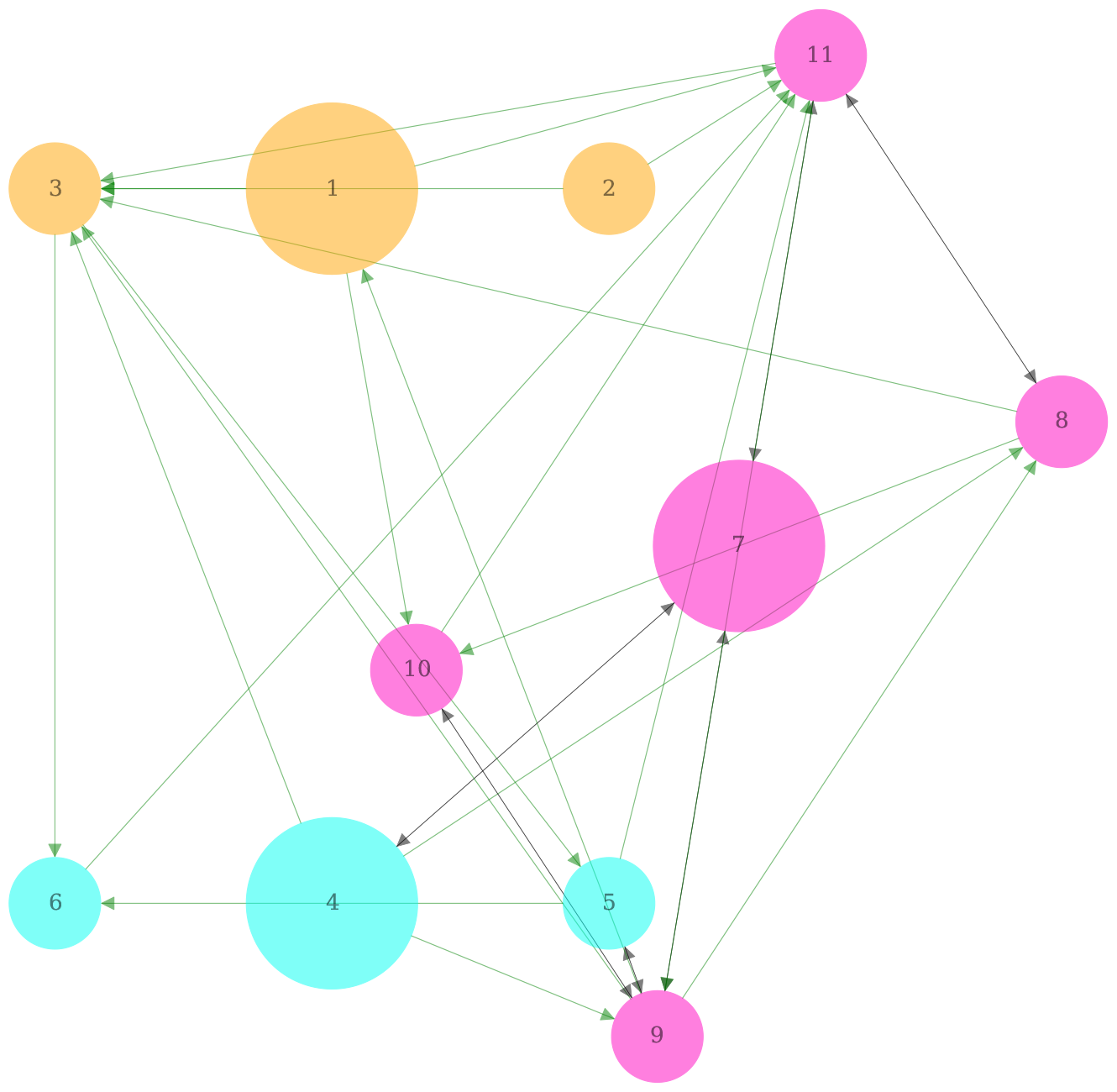
An example of a visualization of the microscopic network.

**Figure (2).**
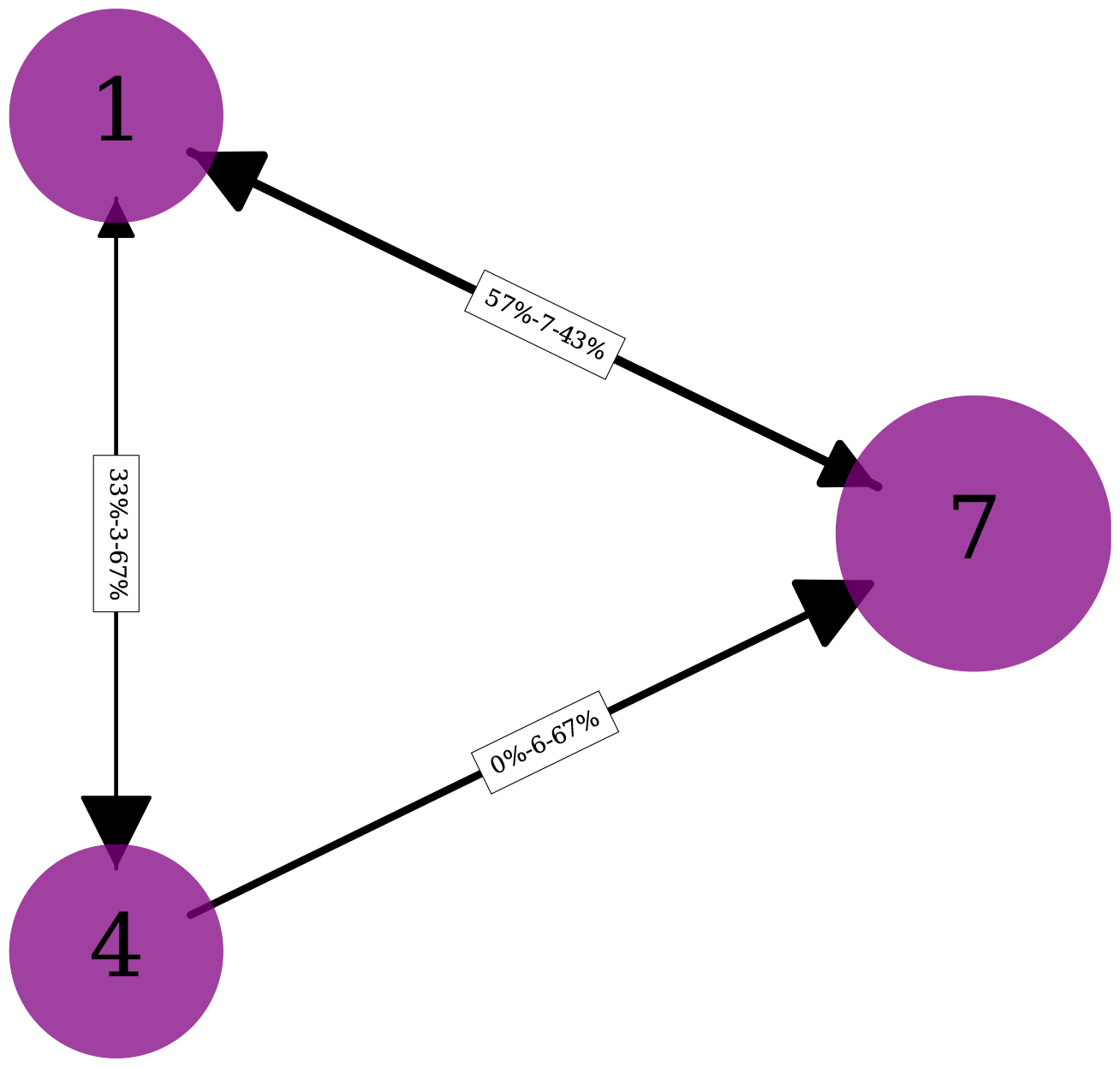
An example of a visualization of the mesoscopic network. Edge labels read as follows: *“% of connections right → left - Total number of connections - % of connections left → right”*.

In the case of the mesoscopic network representation, illustrated in Figure 2, the information on the distributions, encoded in the edges characteristics, is also provided in a form of labels. For example, the edge between the clusters 1 and 2, labeled with the corresponding centroids 1 and 4, has “33%-3-67%” to describe the connectivity distribution. Indeed, the microscopic representation in Figure 1 clarifies this label, noticing that there are 3 directed (green) connections of the members of cluster 1 with those of cluster 2, with 2 directed towards cluster 2, and 1 towards cluster 1.

## 3 Results

### 3.1 Data preprocessing

We estimated transcriptomic fold changes of 157 genes for the LINAC dataset and 152 for the SARRP dataset. Gene expression was measured at 2, 4, 7, 14 and 21 days after irradiation. The estimation was performed based on 3 or 4 replicates. We performed the preprocessing of the raw gene expression data according to the same procedure as presented in Arsenteva et al. (2023). The data is divided by the standard deviation in order to lower relative weight the fold changes with higher uncertainty, and by the fold change norm, defined for a fold change 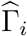 as

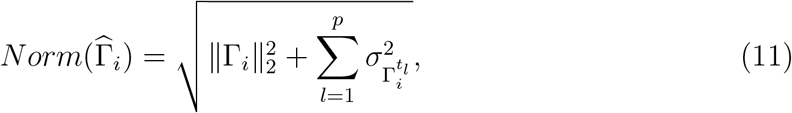

in order to reduce the scale differences between the fold changes. The second step is motivated by the fact that from the radiobiological point of view it is reasonable to ignore the scale differences in expression between genes while performing clustering. For example, if two genes are significantly positively expressed at the same time, but with different strength, we expect them to be in the same cluster, which would be less likely to happen without this transformation.

### 3.2 Choice of the number of clusters

After the fold changes were estimated from the log-transformed data and then preprocessed, we performed clustering coupled with alignment based on the sign-penalized optimal warping dissimilarity matrix computed for both irradiation types. It has been decided to choose 5 clusters produced by k-medoids clustering for these data based on the appearance of clusters expected from the biological point of view. Choosing the number of clusters is a difficult task that often requires considering both mathematical approaches and the prior information from the domain of application (see, for example, Hastie et al. (2009)). Classical selection criteria such as total cost and silhouette score (Rousseeuw, 1987) appear to favor the smallest number of clusters (Supplementary Figure S4). It appears that 5 is the smallest number of clusters that manage to produce well-separated behavior types. We compared clustering of the LINAC fold changes into 4 and 5 groups and concluded that 5 cluster version separates cluster 1 and 3 that are mixed together in cluster 4 of the 4 cluster version (Supplementary Figures S3 and S5). This separation is justified biologically since it is important to distinguish the fold changes that are up-regulated three weeks after irradiation (cluster 3 of 5-cluster version) from those that are up-regulated early but loose the expression by two weeks after irradiation (cluster 1 of 5-cluster version). Such a distinction cannot be ensured merely by the means of having multiple alignment groups, given that having only 5 time points in the dataset the maximal warping step has to be set at 1.

### 3.3 Cluster and network analysis of datasets SARRP and LINAC

#### 3.3.1 Cluster analysis

The five clusters that were obtained are presented in Supplementary Material in Figures S2 (SARRP) and S3 (LINAC). The colorcode and the legend on the plots allows to identify which warp group (warped backward with respect to the centroid, simultaneous with the centroid and warped forward with respect to the centroid) each gene belongs to. Comparing the unaligned and the aligned versions allows to see more clearly how each group has been transformed in order to get aligned with the centroid. The aligned version is the one used for clustering, allowing to identify global behavior types up to a time shift, whereas the unaligned version allows to identify temporal cascades inside every cluster, i.e. the forward-warped predict simultaneous that predict the backward-warped. It has to be noted that these plots only contain the means of preprocessed fold changes, giving a rough idea of the genes’ behavior but can be at times misleading since clustering is performed on full fold changes, containing not only means but all the information on correlations and uncertainties that can be inferred from the replicates.

The clusters are ordered to match between both conditions with respect to the response types represented by each cluster. Indeed, we manage to obtain very similar response types for both conditions: two clusters 1 and 3 characterized by up-regulation, being strongly up-regulated early and late respectively, a generally down-regulated cluster 5 that is roughly symmetric to cluster 1, and two clusters that manifest change of sign, with 2 being up-regulated early and down-regulated late, and 4 doing the opposite. For both irradiation types cluster 2 appears to be much smaller then the others, while cluster 5 contains almost a third of all fold changes. It can be observed that clusters 1 and 3 show much less striking distinction in case of SARRP compared to LINAC. It can also be noted that clusters 4 and 5 appear to have very overall consistent behavior across conditions, which is more visible in the unaligned case rather than aligned, since their centroids belong to different time groups (in case of cluster 4, the LINAC centroid TIMP3 corresponds to earlier expression then the SARRP centroid IL6, and the opposite is observed in case of cluster 5).

#### 3.3.2 Mesoscopic network analysis

The next step of the statistical analysis consists in inferring the transcriptomic fold changes network according to the procedure described in Section 2.3. The mesoscopic network visualization allows in particular to obtain additional distributional information with respect to cluster migrations thanks to leveraging similarities, information that is inaccessible through studying the contingency table alone (Supplementary Material, Table S11). Depending on the goal, one can choose to work with full networks, or a part of it. For instance, mesoscopic graphs of Figure 3 were build based on condition-specific networks, meaning that the connections that form the inter-cluster distribution edges only appear in the corresponding irradiation condition and not the other, which allows to compare two conditions strictly based on their differences. There are multiple edges that appear in both graphs, the most important being the one between 1 (FAS/ACTA2) and 3 (CD44/TAGLN). Given that the networks are condition-specific, it suggests that there is a big number of genes that travel between clusters 1 and 3, which are rather similar in general for both conditions. The most striking difference between the graphs lies in edges that are present/important for one condition and absent/not important for the other. This seems to be particularly the case between clusters 4 (TIMP3/IL6) and 5 (LTBP4/KRT18), the phenomenon that is hard to interpret since it is not clear whether it is similar genes that change clusters, or genes potentially conserving clusters that change their similarity.

**Figure (3).**
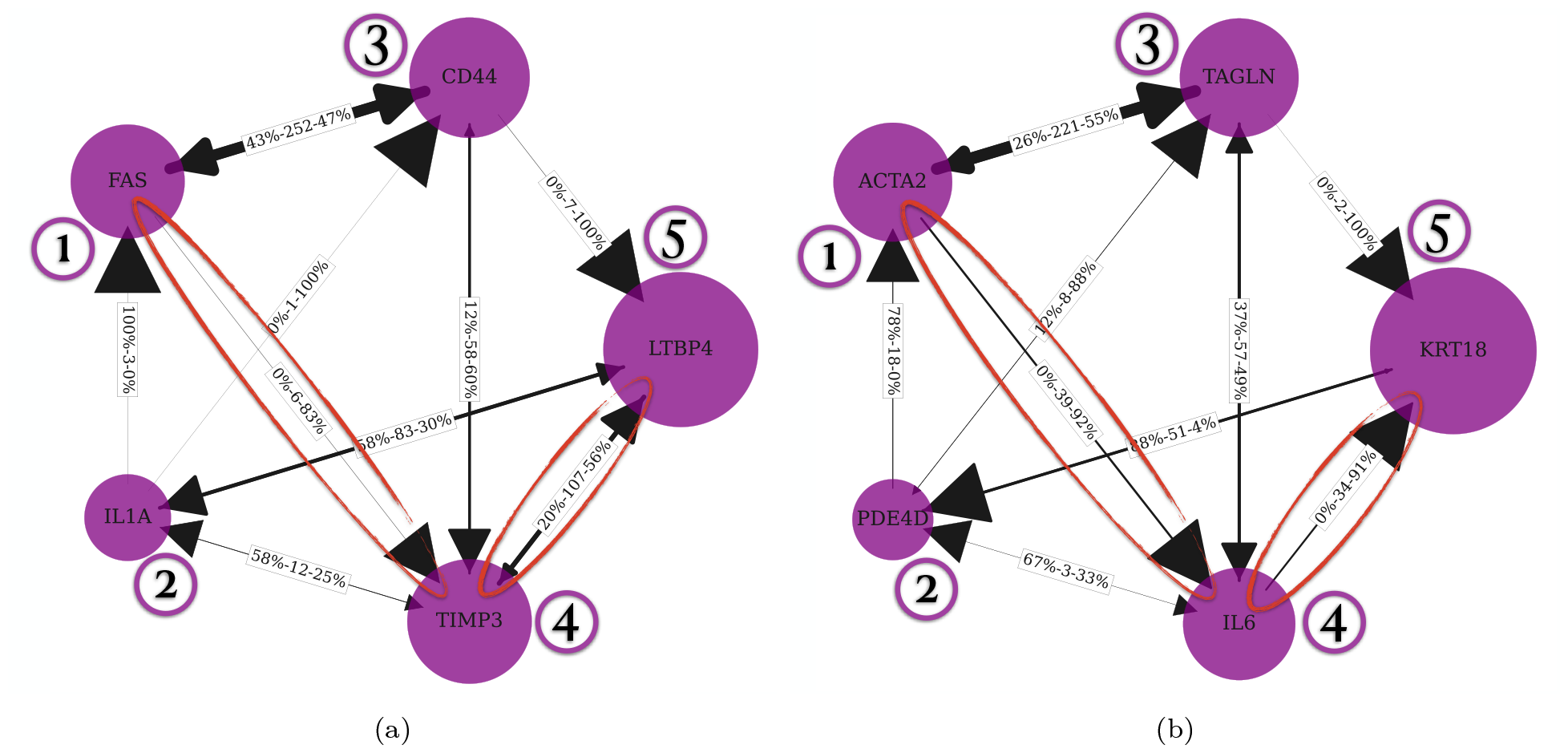
Mesoscopic views of the LINAC-specific (a) and SARRP-specific (b) fold changes network. Nodes represent clusters, labeled with their centroid genes, with node sizes corresponding to respective cluster sizes, and edges summarizing the connections between clusters based on the original adjacency matrix. Edge thickness corresponds to the total number of connections, arrow size to the proportion of predictive connections of the corresponding type.

The situation can be clarified by studying another mesoscopic graph presented in Figure S8 of Supplementary Material, a hybrid representation of two conditions: the genes in each node are those associated with the LINAC clustering, while the network itself (and thus the connections) is the one specific to SARRP. This hybrid graph unsurprisingly demonstrates a bigger overall number of connections since the clustering is not the one natural for the network, and many connections that are otherwise intra-cluster appear here as inter-cluster. In particular, the connection between clusters 4 and 5 is even stronger than that between clusters 1 and 3, which indicates that there is a group of fold changes in LINAC’s clusters 4 and 5 that are extremely similar for both conditions, with connections that disappear in SARRP, which is most likely caused by them becoming intra-cluster connections. All of the above suggests that migrations between clusters 4 and 5 are particularly important in detecting the differences between the two irradiation types.

In order to confirm the significance of the difference between the condition-specific mesoscopic graphs overall and between certain connections in particular, we performed a number of statistical tests on these connections. The tests were applied on the data associated with the mesoscopic graphs presented in Figure 3. First, we performed a Pearson’s Chi-squared test for count data on the edge thickness weights to test their distributional equality. The test is significant, with p-value of 0.0005, we therefore conclude that the number of connections between clusters in the two graphs is statistically different. Next, performed a Barnard exact test on each pair of clusters to compare the significance of the difference between the corresponding numbers of connections for LINAC and SARRP.

For every cluster pair, the test was applied to a 2×2 contingency table, such that the two columns corresponded to LINAC and SARRP, whereas the two rows contained the number of connections between the chosen clusters, and the total number of connection for the dataset in question. The results, presented in Table 1, confirm that the most significantly different connections are indeed those between clusters 1 and 4 as well as clusters 4 and 5. In addition, we repeated the latter test for the mesoscopic networks obtained with different sparsity levels, with the goal of determining to what extent the results obtained based on the mesoscopic graphs are sensitive to the choice of the sparsity level.The obtained p-values are presented in Supplementary Table S12. For 13 sparsity levels between 60% and 90%, we obtained consistently significant differences for the connections of cluster 4 with clusters 1 and 5. Some other connections, for instance those around the smallest cluster 2, are significant for some but not all sparsity levels. This provides even stronger evidence supporting the importance of the role played by cluster 4 and its migrations to clusters 1 and 5 in detecting the differences between the two radiation treatments.

**Table (1).** P-values associated with the results of the Barnard exact test performed on every connection of mesoscopic condition-specific graphs obtained for LINAC and SARRP.

| Link | 1-2 | 1-3 | 1-4 | 1-5 | 2-3 | 2-4 | 2-5 | 3-4 | 3-5 | 4-5 |
| --- | --- | --- | --- | --- | --- | --- | --- | --- | --- | --- |
| P-value | 0.005 | 0.5479 | 0.0006 | 1 | 0.0339 | 0.0682 | 0.1304 | 0.3585 | 0.2193 | 0.0001 |

#### 3.3.3 Microscopic network analysis

The second proposed tool for network visualization sheds additional light on fold changes’ distribution with respect to cluster migration. Figures S9 and S10 of Supplementary Material are two examples of multiple modes of representing a microscopic network with different features. The first figure illustrates the most natural representation mode of the LINAC network, whereas the second figure shows a hybrid network that serves specifically to compare two types of irradiation. For a consistent comparison both are based on the network containing only the links that are shared by both datasets. By comparing the graphs, a few observations can be made that support the conclusions made out of the mesoscopic representations. In particular, it is made clear that approximately a half of LINAC’s cluster 4 becomes a part of cluster 5 for SARRP. It implies that this very group of genes (pink nodes at the cluster 4 position) is responsible for a big number of previously mentionned connections turning intra-cluster. Moreover, cluster 5 is the only cluster whose centroid for SARRP is in cluster 4 for LINAC. However, cluster 5 seems to be much more stable than other clusters overall given its superior size. It can also be noted that clusters 1 and 3 seem to exchange genes mainly between each other, which is consistent with the idea of them being more similar for SARRP than for LINAC.

### 3.4 Enrichment analysis of SARRP and LINAC clusters

From the biological viewpoint, the enrichment analysis of the biological processes for each cluster after radiation at 220 kV (SARRP) has been carried out (Supplementary Figures S11-S15). Cluster 1 is characterized by 2 major functions mainly associated with adhesion and migration process but also with the apoptotic process or the cellular DNA damage response. Cluster 2 has fewer terms than Cluster 1 and is mainly defined by the regulation of signaling pathways such as the pi3kinase pathway which has been described in the literature as involved in the radiation response of endothelial cells (Edwards et al., 2002; Yentrapalli et al., 2013). Cluster 3 is characterized by TGFbeta and SMAD family related terms, which have been previously described in the literature on the endothelial vascular response and a glycosylation related term emerged from the enrichment analysis of cluster 3 (Milliat et al., 2006; Jaillet et al., 2017; Ladaigue et al., 2022). Cluster 4 is related to the function of the chemoattraction and cell-cell interaction with the immune system. These major features can be compared with the results for cluster 1 with respect to the term of adhesion. Moreover, in cluster 4 we see appear terms related to the control of the apoptotic process, a feature that also appears in the cluster 1. Cluster 4 is also linked to coagulation processes and fibrinolysis, described as a mark of radiation-induced endothelial response (Milliat et al., 2008). Lastly, cluster 5 is characterized by the activation of phosphorylation signaling pathways such as SMADs or the NFKB pathway. This can also be related to the enrichment of TGF feta in cluster3. A protein glycosylation term, previously captured in cluster 3, can also be found in cluster 5, which is in accordance with the results published in the literature showing an impact of glycosylation process in the radiation response of endothelial cells (Jaillet et al., 2017; Ladaigue et al., 2022). The results of the enrichment analysis coupled with the cluster and network analyses are illustrated in Figure 4.

**Figure (4).**
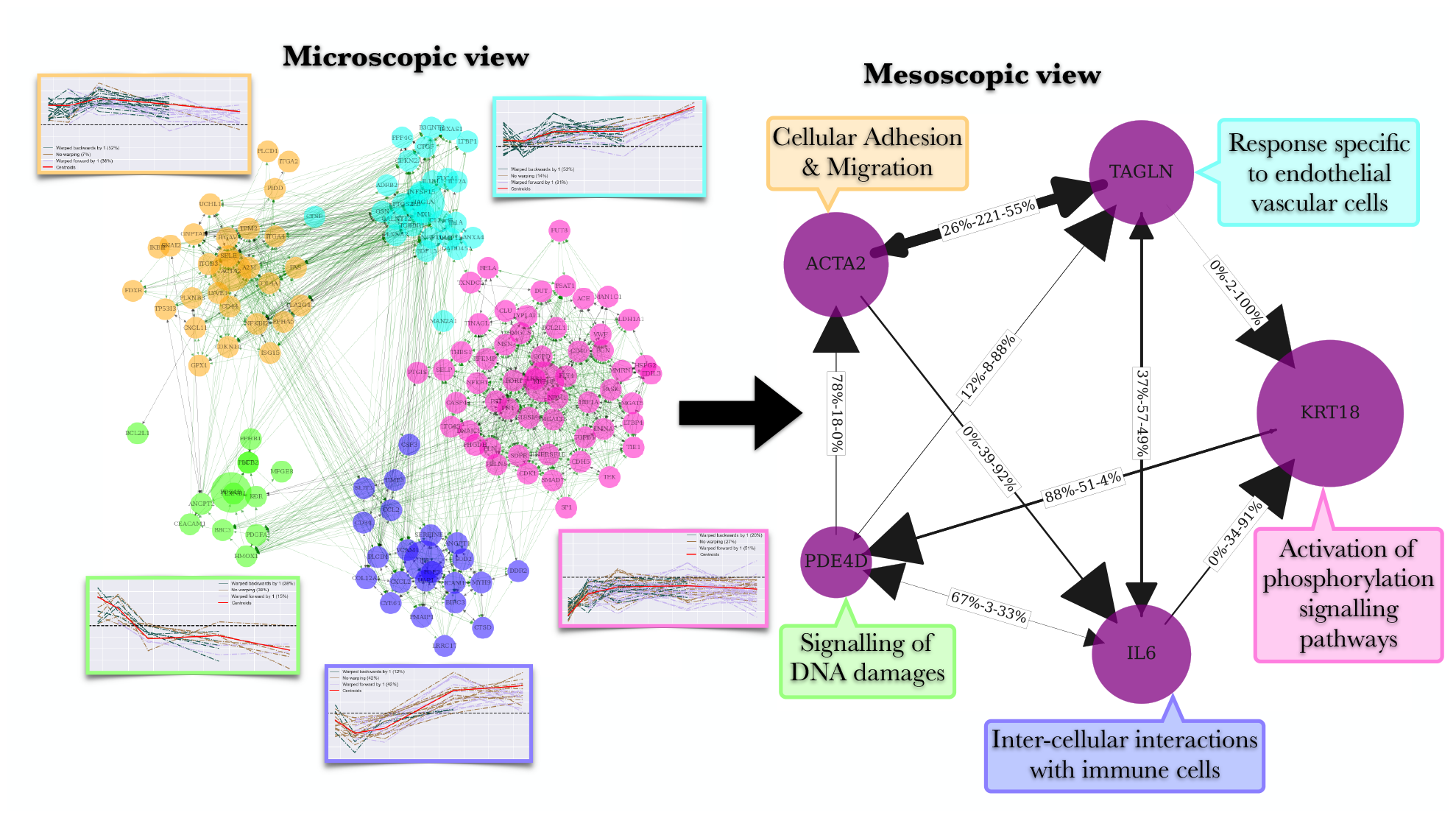
Summary of the cluster and network analyses performed on the SARRP dataset with the key biological functions identified as a result of the enrichment analysis.

The same analysis has been performed on the dataset obtained after irradiation at 4 MV (LINAC). The comparison of deregulated genes after radiation in the two irradiation conditions is illustrated through the Venn diagrams in Supplementary Figures S16-S20. The enrichment analysis of biological processes (Supplementary Figures S21-S25) reveals that irradiation at 4 MV has an impact on biological processes on all clusters. In particular, there are senescence-related terms that appear for 4 MV but are absent for 220 kV. This is consistent with the previously published results showing that cellular senescence is more important at 4 MV than at 220 kV (Paget et al., 2019), this illustrating the robustness of the analytical methodological approach implemented in this work. Moreover, these results illustrate by an innovative approach that the RBE of photons at 4 MV compared to 220 kV is not one.

The mathematical model also predicts a particular affinity for clusters 1 and 4 after irradiation at 220 kV and clusters 4 and 5 after irradiation at 4 MV suggesting that the terms appear in these various clusters may potentially explain the differences in response to the 2 energies. Combining the enrichment analyses of clusters 1 and 4 for 220 kV and clusters 4 and 5 for 4 MV, the terms that were globally identified focus mainly on cell adhesion and chemotaxis, suggesting a global energy-dependent effect on these inflammation-related parameters. It has already been shown that radiation response of endothelial cells after 4MV compared with 220 kV is characterized by more senescence and more inflammatory induced response with upregulation of IL6 and IL8 higher at 4 MV than at 220 kV (Paget et al., 2019). These results reinforce the idea that the physical dose in Gray is not sufficient to predict a biological effect and by extrapolation a risk. Our results open biological hypotheses concerning the impact of radiation used in the medical field and in particular radiotherapy on both tumors and healthy tissues.

## 4 Conclusion

In this work we performed the analysis of two in vitro datasets corresponding to two different radiation modalities. We proposed new tools for network-based key features visualization, designed for complex omics datasets. The tools are based on the statistical framework performing estimation and subsequent clustering jointly with alignment of the omic fold changes. We identified the major response types and the associated temporal cascades of genes. The proposed tools provided easy results interpretability and subsequent hypotheses generation, the pertinence of these results has been supported by the enrichment analysis. Applied to datasets associated with a variety of treatments by radiotherapy, our methodology can provide insight on the mechanisms behind the response to these treatments, bringing us one step closer to minimizing the potential adverse effects.

## 6 Author notes

PA, HC and MAB contributed to the conception and design of the study. PA constructed the analysis pipeline and the Python package. PA, HC, MAB interpreted the mathematical results. FM, OG, VP, GT, MDS collected the experimental data and biological interpretation. PA, HC, MAB, FM, OG wrote the first manuscript draft. All authors contributed to manuscript revision, read, and approved the submitted version.

