## Supplementary tables and figures for "Comparing cellular response to two radiation treatments based on key features visualization"

| Protein/ | Full name | Gene probe |
|---|---|---|
| AKAP12 | A kinase (PRKA) anchor protein 12 | Hs00374507_m1 |
| ALDH1A1 | aldehyde dehydrogenase 1 family, member A1 | Hs00946916_m1 |
| ANXA4 | annexin A4 | Hs00984874_m1 |
| BGN | biglycan | Hs00156076_m1 |
| C12orf5 | chromosome 12 open reading frame 5 | Hs00608644_m1 |
| CAP1 | CAP, adenylate cyclase-associated protein 1 (yeast) | Hs00255173_m1 |
| CDK1 | cyclin-dependent kinase 1 | Hs00938777_m1 |
| CLU | clusterin | Hs00156548_m1 |
| COL12A1 | collagen, type XII, alpha 1 | Hs00189184_m1 |
| CTGF | connective tissue growth factor | Hs00170014_m1 |
| CTSB | cathepsin B | Hs00947433_m1 |
| CTSD | cathepsin D | Hs00157205_m1 |
| CYR61 | cysteine-rich, angiogenic inducer, 61 | Hs00155479_m1 |
| DNAJC9 | DnaJ (Hsp40) homolog, subfamily C, member 9 | Hs00750059_s1 |
| DUT | deoxyuridine triphosphatase | Hs00798995_s1 |
| EDIL3 | EGF-like repeats and discoidin I-like domains 3 | Hs00964112_m1 |
| EFEMP1 | EGF containing fibulin-like extracellular matrix protein 1 | Hs00244575_m1 |
| ELN | elastin | Hs00355783_m1 |
| FBLN1 | fibulin 1 | Hs00972609_m1 |
| FDXR | ferredoxin reductase | Hs01031617_m1 |
| FN1 | fibronectin 1 | Hs00365052_m1 |
| G6PD | glucose-6-phosphate dehydrogenase | Hs00166169_m1 |
| GDF15 | growth differentiation factor 15 | Hs00171132_m1 |
| GPX1 | glutathione peroxidase 1 | Hs01028922_g1 |
| GSN | gelsolin | Hs00609272_m1 |
| HMGCS1 | 3-hydroxy-3-methylglutaryl-CoA synthase 1 (soluble) | Hs00940429_m1 |
| HSPG2 | heparan sulfate proteoglycan 2 | Hs00194179_m1 |
| IKBIP | IKBKB interacting protein | Hs01096287_m1 |
| ISG15 | ISG15 ubiquitin-like modifier | Hs00192713_m1 |
| ITGA2 | integrin, alpha 2 (CD49B, alpha 2 subunit of VLA-2 receptor) | Hs00158127_m1 |
| KRT18 | keratin 18 | Hs02827483_g1 |
| LMNA | lamin A/C | Hs00153462_m1 |
| LYPLAL1 | lysophospholipase-like 1 | Hs00371117_m1 |
| MFGE8 | milk fat globule-EGF factor 8 protein | Hs00170712_m1 |
| MMRN1 | multimerin 1 | Hs00201182_m1 |
| MSN | moesin | Hs00741306_mH |
| MX1 | myxovirus (influenza virus) resistance 1, interferon-inducible protein p78 (mouse) | Hs00895608_m1 |
| MYH9 | myosin, heavy chain 9, non-muscle | Hs00159522_m1 |
| NPM1 | nucleophosmin (nucleolar phosphoprotein B23, numatrin) | Hs02339479_g1 |
| PHGDH | phosphoglycerate dehydrogenase | Hs00198333_m1 |
| PPP4C | protein phosphatase 4, catalytic subunit | Hs00427262_m1 |
| PSAT1 | phosphoserine aminotransferase 1 | Hs00253548_m1 |
| SDPR | serum deprivation response | Hs00190538_m1 |
| SERPINE1 | serpin peptidase inhibitor, clade E (nexin, plasminogen activator inhibitor type 1), member 1 | Hs01126606_m1 |
| SOD2 | superoxide dismutase 2, mitochondrial | Hs00167309_m1 |
| TAGLN | transgelin | Hs00162558_m1 |
| THBS1 | thrombospondin 1 | Hs00962908_m1 |
| TINAGL1 | tubulointerstitial nephritis antigen-like 1 | Hs01115879_m1 |
| TP53I3 | tumor protein p53 inducible protein 3 | Hs00936520_m1 |
| TPM2 | tropomyosin 2 (beta) | Hs00268540_m1 |
| TXNDC5 | thioredoxin domain containing 5 (endoplasmic reticulum) | Hs01046713_m1 |
| UAP1 | UDP-N-acteylglucosamine pyrophosphorylase 1 | Hs00268394_m1 |
| UCHL1 | ubiquitin carboxyl-terminal esterase L1 (ubiquitin thiolesterase) | Hs00985157_m1 |
| VWF | von Willebrand factor | Hs01109446_m1 |
| A2M | alpha-2-macroglobulin | Hs00929971_m1 |
| ACE | angiotensin I converting enzyme | Hs00174179_m1 |
| ADRB2 | adrenoceptor beta 2, surface | Hs00240532_s1 |
| ANGPT1 | angiopoietin 1 | Hs00375822_m1 |
| ANGPT2 | angiopoietin 2 | Hs01048042_m1 |
| ANGPTL1 | angiopoietin-like 1 | Hs00559786_m1 |
| ANGPTL4 | angiopoietin-like 4 | Hs01101127_m1 |
| BAI1 | brain-specific angiogenesis inhibitor 1 | Hs00181777_m1 |
| BBC3 | BCL2 binding component 3 | Hs00248075_m1 |

Table (S1) Genes included in the custom TaqMan Low-Density Array (TLDA) to measure the expression levels of 192 mRNAs involved in the response of HUVECs to ionizing radiation.

| Protein/Gene ID | Full name | Gene probe |
|---|---|---|
| BCL2A1 | BCL2-related protein A1 | Hs00187845_m1 |
| BCL2L1 | BCL2-like 1 | Hs00236329_m1 |
| BCL2L11 | BCL2-like 11 (apoptosis facilitator) | Hs01076940_m1 |
| BDKRB2 | bradykinin receptor B2 | Hs00176121_m1 |
| BIRC3 | baculoviral IAP repeat containing 3 | Hs00154109_m1 |
| BIRC5 | baculoviral IAP repeat containing 5 | Hs00978503_m1 |
| BMPR1B | bone morphogenetic protein receptor, type IB | Hs01010965_m1 |
| C3 | complement component 3 | Hs00163811_m1 |
| CASP4 | caspase 4, apoptosis-related cysteine peptidase | Hs01031951_m1 |
| CCL5 | chemokine (C-C motif) ligand 5 | Hs00982282_m1 |
| CCR4 | chemokine (C-C motif) receptor 4 | Hs99999919_m1 |
| CD34 | CD34 molecule | Hs00990732_m1 |
| CD44 | CD44 molecule (Indian blood group) | Hs01075861_m1 |
| CEACAM1 | carcinoembryonic antigen-related cell adhesion molecule 1 (biliary glycoprotein) | Hs00989786_m1 |
| COL15A1 | collagen, type XV, alpha 1 | Hs00266332_m1 |
| COL4A1 | collagen, type IV, alpha 1 | Hs00266237_m1 |
| CSF2 | colony stimulating factor 2 (granulocyte-macrophage) | Hs00929873_m1 |
| CXCL10 | chemokine (C-X-C motif) ligand 10 | Hs00171042_m1 |
| CXCL11 | chemokine (C-X-C motif) ligand 11 | Hs00171138_m1 |
| CXCL2 | chemokine (C-X-C motif) ligand 2 | Hs00601975_m1 |
| DDR2 | discoidin domain receptor tyrosine kinase 2 | Hs01025953_m1 |
| EPHA5 | EPH receptor A5 | Hs00300724_m1 |
| EPHB1 | EPH receptor B1 | Hs01057849_m1 |
| FAS | Fas cell surface death receptor | Hs00236330_m1 |
| FBLN5 | fibulin 5 | Hs00197064_m1 |
| FGF1 | fibroblast growth factor 1 (acidic) | Hs01092738_m1 |
| FGF2 | fibroblast growth factor 2 (basic) | Hs00266645_m1 |
| FLT4 | fms-related tyrosine kinase 4 | Hs01047677_m1 |
| FST | follistatin | Hs00246256_m1 |
| GFRA1 | GDNF family receptor alpha 1 | Hs00237133_m1 |
| HLA-DRB1 | major histocompatibility complex, class II, DR beta 1 | Hs99999917_m1 |
| HMOX1 | heme oxygenase (decycling) 1 | Hs01110250_m1 |
| ICAM1 | intercellular adhesion molecule 1 | Hs00164932_m1 |
| IL12A | interleukin 12A (natural killer cell stimulatory factor 1, cytotoxic lymphocyte maturation factor 1, p35) | Hs01073447_m1 |
| IL1A | interleukin 1, alpha | Hs00174092_m1 |
| IL1RL1 | interleukin 1 receptor-like 1 | Hs00545033_m1 |
| ITGA4 | integrin, alpha 4 (antigen CD49D, alpha 4 subunit of VLA-4 receptor) | Hs00168433_m1 |
| ITGAV | integrin, alpha V | Hs00233808_m1 |
| ITGB3 | integrin, beta 3 (platelet glycoprotein IIIa, antigen CD61) | Hs01001469_m1 |
| KDR | kinase insert domain receptor (a type III receptor tyrosine kinase) | Hs00911700_m1 |
| KIT | v-kit Hardy-Zuckerman 4 feline sarcoma viral oncogene homolog | Hs00174029_m1 |
| LRDD | p53-induced death domain protein | Hs00388035_m1 |
| LRR1 | leucine rich repeat protein 1 | Hs00381120_m1 |
| LRRC17 | leucine rich repeat containing 17 | Hs00180581_m1 |
| LTB | lymphotoxin beta (TNF superfamily, member 3) | Hs00242737_m1 |
| LTBP1 | latent transforming growth factor beta binding protein 1 | Hs00386448_m1 |
| LTBP4 | latent transforming growth factor beta binding protein 4 | Hs00186025_m1 |
| LTC4S | leukotriene C4 synthase | Hs00168529_m1 |
| LYVE1 | lymphatic vessel endothelial hyaluronan receptor 1 | Hs00272659_m1 |
| NFKBIZ | nuclear factor of kappa light polypeptide gene enhancer in B-cells inhibitor, zeta | Hs00230071_m1 |
| NTRK2 | neurotrophic tyrosine kinase, receptor, type 2 | Hs00178811_m1 |
| PASK | PAS domain containing serine/threonine kinase | Hs00209470_m1 |
| PDE4D | phosphodiesterase 4D, cAMP-specific | Hs01579625_m1 |
| PDGFRB | platelet-derived growth factor receptor, beta polypeptide | Hs01019589_m1 |
| PLA2G4C | phospholipase A2, group IVC (cytosolic, calcium-independent) | Hs01003743_m1 |
| PLA2G5 | phospholipase A2, group V | Hs00173472_m1 |
| PLCB2 | phospholipase C, beta 2 | Hs01080542_m1 |
| PLCB4 | phospholipase C, beta 4 | Hs01075770_m1 |
| PLCD1 | phospholipase C, delta 1 | Hs00979908_m1 |
| PLXNA3 | plexin A3 | Hs00250178_m1 |
| PLXNB1 | plexin B1 | Hs00963507_m1 |
| PLXNB3 | plexin B3 | Hs00193613_m1 |
| PMAIP1 | phorbol-12-myristate-13-acetate-induced protein 1 | Hs00560402_m1 |

Table (S1) Genes included in the custom TaqMan Low-Density Array (TLDA) to measure the expression levels of 192 mRNAs involved in the response of HUVECs to ionizing radiation (continued).

| Protein/Gene ID | Full name | Gene probe |
|---|---|---|
| PTGIS | prostaglandin I2 (prostacyclin) synthase | Hs00919949_m1 |
| PTGS2 | prostaglandin-endoperoxide synthase 2 (prostaglandin G/H synthase and cyclooxygenase) | Hs00153133_m1 |
| ROR1 | receptor tyrosine kinase-like orphan receptor 1 | Hs00938677_m1 |
| SELE | selectin E | Hs00950401_m1 |
| SELP | selectin P (granule membrane protein 140kDa, antigen CD62) | Hs00927900_m1 |
| SLIT3 | slit homolog 3 (Drosophila) | Hs00171524_m1 |
| SMAD7 | SMAD family member 7 | Hs00998193_m1 |
| TBXAS1 | thromboxane A synthase 1 (platelet) | Hs01022706_m1 |
| TGFBR1 | transforming growth factor, beta receptor 1 | Hs00610320_m1 |
| TIMP3 | TIMP metallopeptidase inhibitor 3 | Hs00165949_m1 |
| TNFRSF1B | tumor necrosis factor receptor superfamily, member 1B | Hs00961749_m1 |
| TNFSF15 | tumor necrosis factor (ligand) superfamily, member 15 | Hs00270802_s1 |
| VCAM1 | vascular cell adhesion molecule 1 | Hs01003372_m1 |
| 18S | 18S rRNA | Hs99999901_s1 |
| ACTB | actin, beta | Hs99999903_m1 |
| MAPK8 | mitogen-activated protein kinase 8 | Hs00177083_m1 |
| PGK1 | phosphoglycerate kinase 1 | Hs99999906_m1 |
| SKI | v-ski avian sarcoma viral oncogene homolog | Hs00161707_m1 |
| CD40 | CD40 molecule, TNF receptor superfamily member 5 | Hs00386848_m1 |
| CXCL12 | chemokine (C-X-C motif) ligand 12 | Hs00171022_m1 |
| HIF1A | hypoxia inducible factor 1, alpha subunit (basic helix-loop-helix transcription factor) | Hs00153153_m1 |
| NFKB1 | nuclear factor of kappa light polypeptide gene enhancer in B-cells 1 | Hs00765730_m1 |
| PDGFA | platelet-derived growth factor alpha polypeptide | Hs00964426_m1 |
| RELA | v-rel avian reticuloendotheliosis viral oncogene homolog A | Hs00153294_m1 |
| SP1 | Sp1 transcription factor | Hs00916521_m1 |
| TGFB1 | transforming growth factor, beta 1 | Hs00998133_m1 |
| ACTA2 | alpha smooth muscle actin | Hs00909449_m1 |
| CDH5 | VE-Cadherin | Hs00901463_m1 |
| HEY2 | Hairy/enhancer-of-split related with YRPW motif protein 2 | Hs00232622_m1 |
| SNAIL2 | Snail family zing finger 2 (slug) | Hs00950344_m1 |
| SPP1 | Osteopontin | Hs00959010_m1 |
| TEK | tyrosine kinase with immunoglobulin-like and EGF-like domains 2 | Hs00945146_m1 |
| TIE1 | tyrosine kinase with immunoglobulin-like and EGF-like domains 1 | Hs00892696_m1 |
| TNC | Tenascine C | Hs01115665_m1 |
| VTN | Vitronectine | Hs00169863_m1 |
| CCL2 | chemokine (C-C motif) ligand 2 | Hs00234140_m1 |
| CCL4 | chemokine (C-C motif) ligand 4 | Hs01031494_m1 |
| CSF3 | colony stimulating factor 3 (granulocyte) | Hs99999083_m1 |
| IFNG | interferon, gamma | Hs00989291_m1 |
| IL10 | interleukin 10 | Hs00961622_m1 |
| IL17A | interleukin 17A | Hs00174383_m1 |
| IL1B | interleukin 1, beta | Hs01555410_m1 |
| IL2 | interleukin 2 | Hs00174114_m1 |
| IL4 | interleukin 4 | Hs00174122_m1 |
| IL6 | interleukin 6 (interferon, beta 2) | Hs00985639_m1 |
| IL8 | interleukin 8 | Hs00174103_m1 |
| TNF | tumor necrosis factor | Hs99999043_m1 |
| CDKN1A | cyclin-dependent kinase inhibitor 1A (p21, Cip1) | Hs00355782_m1 |
| CDKN2A | cyclin-dependent kinase inhibitor 2A | Hs00923894_m1 |
| GADD45A | growth arrest and DNA-damage-inducible, alpha | Hs00169255_m1 |
| FUCA1 | fucosidase, alpha-L- 1, tissue | Hs00609173_m1 |
| FUT8 | fucosyltransferase 8 (alpha (1,6) fucosyltransferase) | Hs00189535_m1 |
| GALNT12 | UDP-N-acetyl-alpha-D-galactosamine:polypeptide N-acetylgalactosaminyltransferase 12 (GalNAc-T12) | Hs00226436_m1 |
| GALNT14 | UDP-N-acetyl-alpha-D-galactosamine:polypeptide N-acetylgalactosaminyltransferase 14 (GalNAc-T14) | Hs00226180_m1 |
| GCNT3 | glucosaminyl (N-acetyl) transferase 3, mucin type | Hs00953355_m1 |
| GCNT4 | glucosaminyl (N-acetyl) transferase 4, core 2 | Hs00275464_s1 |
| GNPTAB | N-acetylglucosamine-1-phosphate transferase, alpha and beta subunits | Hs00225647_m1 |
| MAN1C1 | mannosidase, alpha, class 1C, member 1 | Hs00220595_m1 |
| MAN2A1 | mannosidase, alpha, class 2A, member 1 | Hs00159007_m1 |
| MGAT5 | mannosyl (alpha-1,6-)-glycoprotein beta-1,6-N-acetyl-glucosaminyltransferase | Hs01073268_m1 |
| ST8SIA6 | ST8 alpha-N-acetyl-neuraminide alpha-2,8-sialyltransferase 6 | Hs02341873_m1 |
| B3GNT2 | UDP-GlcNAc:betaGal beta-1,3-N-acetylglucosaminyltransferase 2 | Hs00198128_m1 |
| B4GALT5 | UDP-Gal:betaGlcNAc beta 1,4- galactosyltransferase, polypeptide 5 | Hs00941041_m1 |

Table (S1) Genes included in the custom TaqMan Low-Density Array (TLDA) to measure the expression levels of 192 mRNAs involved in the response of HUVECs to ionizing radiation (continued).

| Protein/ Gene ID | Full name |
|---|---|
| AKAP12 | A kinase (PRKA) anchor protein 12 |
| ALDH1A1 | Aldehyde dehydrogenase 1 family, member A1 |
| ANXA4 | Annexin A4 |
| BGN | Biglycan |
| C12orf5 | Chromosome 12 open reading frame 5 |
| CAP1 | CAP, adenylate cyclase-associated protein 1 (yeast) |
| CDK1 | Cyclin-dependent kinase 1 |
| CLU | Clusterin |
| COL12A1 | Collagen, type XII, alpha 1 |
| CTGF | Connective tissue growth factor |
| CTSB | Cathepsin B |
| CTSD | Cathepsin D |
| CYR61 | Cysteine-rich, angiogenic inducer, 61 |
| DNAJC9 | Dnaj (hsp40) homolog, subfamily C, member 9 |
| DUT | Deoxyuridine triphosphatase |
| EDIL3 | Egf-like repeats and discoidin i-like domains 3 |
| EFEMP1 | EGF containing fibulin-like extracellular matrix protein 1 |
| ELN | Elastin |
| FBLN1 | Fibulin 1 |
| FDXR | Ferredoxin reductase |
| FN1 | Fibronectin 1 |
| G6PD | Glucose-6-phosphate dehydrogenase |
| GDF15 | Growth differentiation factor 15 |
| GPX1 | Glutathione peroxidase 1 |
| GSN | Gelsolin |
| HIST1H4A | Histone cluster 1, h4a |
| HMGCS1 | 3-hydroxy-3-methylglutaryl-coa synthase 1 (soluble) |
| HSPG2 | Heparan sulfate proteoglycan 2 |
| IKBIP | IKBKB interacting protein |
| ISG15 | ISG15 ubiquitin-like modifier |
| ITGA2 | Integrin, alpha 2 (CD49B, alpha 2 subunit of VLA-2 receptor) |
| KPNA4 | Karyopherin alpha 4 (importin alpha 3) |
| KRT18 | Keratin 18 |
| KRT7 | Keratin 7 |
| LAP3 | Leucine aminopeptidase 3 |
| LMNA | Lamin A/C |
| LYPLAL1 | Lysophospholipase-like 1 |
| MFGE8 | Milk fat globule-egf factor 8 protein |
| MMRN1 | Multimerin 1 |
| MSN | Moesin |
| MX1 | Myxovirus (influenza virus) resistance 1, interferon-inducible protein p78 (mouse) |
| MYH9 | Myosin, heavy chain 9, non-muscle |
| NPM1 | Nucleophosmin (nucleolar phosphoprotein B23, numatrin) |
| PHGDH | Phosphoglycerate dehydrogenase |
| PPP4C | Protein phosphatase 4, catalytic subunit |
| PRPF19 | Pre-mrna processing factor 19 |
| PSAT1 | Phosphoserine aminotransferase 1 |
| SDPR | Serum deprivation response |
| SERPINE1 | Serpin peptidase inhibitor, clade E (nexin, plasminogen activator inhibitor type 1), member 1 |
| SOD2 | Superoxide dismutase 2, mitochondrial |
| TAGLN | Transgelin |
| THBS1 | Thrombospondin 1 |
| TINAGL1 | Tubulointerstitial nephritis antigen-like 1 |
| TP53I3 | Tumor protein p53 inducible protein 3 |
| TPM2 | Tropomyosin 2 (beta) |
| TXNDC5 | Thioredoxin domain containing 5 (endoplasmic reticulum) |
| UAP1 | Udp-n-acteylglucosamine pyrophosphorylase 1 |
| UCHL1 | Ubiquitin carboxyl-terminal esterase L1 (ubiquitin thiolesterase) |
| VWF | Von willebrand factor |

Table (S2) Genes selected following proteomic analysis to generate the transcriptomic signature of irradiation.

| Protein/ Gene ID | Full name |
|---|---|
| A2M | Alpha-2-macroglobulin |
| ACE | Angiotensin I Converting Enzyme |
| ADRB2 | Adrenoceptor Beta 2, Surface |
| ANGPT1 | Angiopoietin 1 |
| ANGPT2 | Angiopoietin 2 |
| ANGPTL1 | Angiopoietin-like 1 |
| ANGPTL4 | Angiopoietin-like 4 |
| BAI1 | Brain-specific Angiogenesis Inhibitor 1 |
| BBC3 | BCL2 Binding Component 3 |
| BCL2A1 | Bcl2-related Protein A1 |
| BCL2L1 | Bcl2-like 1 |
| BCL2L11 | Bcl2-like 11 (Apoptosis Facilitator) |
| BDKRB2 | Bradykinin Receptor B2 |
| BIRC3 | Baculoviral IAP Repeat Containing 3 |
| BIRC5 | Baculoviral IAP Repeat Containing 5 |
| BMPR1B | Bone Morphogenetic Protein Receptor, Type IB |
| C3 | Complement Component 3 |
| CASP4 | Caspase 4, Apoptosis-related Cysteine Peptidase |
| CCL5 | Chemokine (C-C Motif) Ligand 5 |
| CCR4 | Chemokine (C-C Motif) Receptor 4 |
| CD34 | CD34 Molecule |
| CD40 | CD40 Molecule, TNF Receptor Superfamily Member 5 |
| CD44 | CD44 Molecule (Indian Blood Group) |
| CEACAM1 | Carcinoembryonic Antigen-related Cell Adhesion Molecule 1 (Biliary Glycoprotein) |
| COL15A1 | Collagen, Type XV, Alpha 1 |
| COL4A1 | Collagen, Type IV, Alpha 1 |
| CSF2 | Colony Stimulating Factor 2 (Granulocyte-macrophage) |
| CSF3 | Colony Stimulating Factor 3 (Granulocyte) |
| CXCL10 | Chemokine (C-X-C Motif) Ligand 10 |
| CXCL11 | Chemokine (C-X-C Motif) Ligand 11 |
| CXCL12 | Chemokine (C-X-C Motif) Ligand 12 |
| CXCL2 | Chemokine (C-X-C Motif) Ligand 2 |
| DDR2 | Discoidin Domain Receptor Tyrosine Kinase 2 |
| EDIL3 | Egf-like Repeats And Discoidin I-like Domains 3 |
| EPHA5 | EPH Receptor A5 |
| EPHB1 | EPH Receptor B1 |
| FAS | Fas Cell Surface Death Receptor |
| FBLN5 | Fibulin 5 |
| FGF1 | Fibroblast Growth Factor 1 (Acidic) |
| FGF2 | Fibroblast Growth Factor 2 (Basic) |
| FLT4 | Fms-related Tyrosine Kinase 4 |
| FN1 | Fibronectin 1 |
| FST | Follistatin |
| GFRA1 | GDNF Family Receptor Alpha 1 |
| HLA-DRB1 | Major Histocompatibility Complex, Class II, DR Beta 1 |
| HMOX1 | Heme Oxygenase (Decycling) 1 |
| ICAM1 | Intercellular Adhesion Molecule 1 |
| IL12A | Interleukin 12A (Natural Killer Cell Stimulatory Factor 1, Cytotoxic Lymphocyte Maturation Factor 1, P35) |
| IL1A | Interleukin 1, Alpha |
| IL1B | Interleukin 1, Beta |
| IL1RL1 | Interleukin 1 Receptor-like 1 |

Table (S3) Genes selected following gene expression analysis to generate the transcriptomic signature of irradiation.

| Protein/ Gene ID | Full name |
|---|---|
| IL6 | Interleukin 6 (Interferon, Beta 2) |
| IL8 | Interleukin 8 |
| ITGA4 | Integrin, Alpha 4 (Antigen CD49D, Alpha 4 Subunit Of VLA-4 Receptor) |
| ITGAV | Integrin, Alpha V |
| ITGB3 | Integrin, Beta 3 (Platelet Glycoprotein Iiia, Antigen CD61) |
| KDR | Kinase Insert Domain Receptor (A Type III Receptor Tyrosine Kinase) |
| KIT | V-kit Hardy-zuckerman 4 Feline Sarcoma Viral Oncogene Homolog |
| LRDD | P53-induced Death Domain Protein |
| LRR1 | Leucine Rich Repeat Protein 1 |
| LRRC17 | Leucine Rich Repeat Containing 17 |
| LTB | Lymphotoxin Beta (TNF Superfamily, Member 3) |
| LTBP1 | Latent Transforming Growth Factor Beta Binding Protein 1 |
| LTBP4 | Latent Transforming Growth Factor Beta Binding Protein 4 |
| LTC4S | Leukotriene C4 Synthase |
| LYVE1 | Lymphatic Vessel Endothelial Hyaluronan Receptor 1 |
| NFKBIZ | Nuclear Factor Of Kappa Light Polypeptide Gene Enhancer In B-cells Inhibitor, Zeta |
| NTRK2 | Neurotrophic Tyrosine Kinase, Receptor, Type 2 |
| PASK | PAS Domain Containing Serine/Threonine Kinase |
| PDE4D | Phosphodiesterase 4D, Camp-specific |
| PDGFRB | Platelet-derived Growth Factor Receptor, Beta Polypeptide |
| PLA2G4C | Phospholipase A2, Group IVC (Cytosolic, Calcium-independent) |
| PLA2G5 | Phospholipase A2, Group V |
| PLCB2 | Phospholipase C, Beta 2 |
| PLCB4 | Phospholipase C, Beta 4 |
| PLCD1 | Phospholipase C, Delta 1 |
| PLXNA3 | Plexin A3 |
| PLXNB1 | Plexin B1 |
| PLXNB3 | Plexin B3 |
| PMAIP1 | Phorbol-12-myristate-13-acetate-induced Protein 1 |
| PTGIS | Prostaglandin I2 (Prostacyclin) Synthase |
| PTGS2 | Prostaglandin-endoperoxide Synthase 2 (Prostaglandin G/H Synthase And Cyclooxygenase) |
| ROR1 | Receptor Tyrosine Kinase-like Orphan Receptor 1 |
| SELE | Selectin E |
| SELP | Selectin P (Granule Membrane Protein 140kda, Antigen CD62) |
| SLIT3 | Slit Homolog 3 (Drosophila) |
| SMAD7 | SMAD Family Member 7 |
| TBXAS1 | Thromboxane A Synthase 1 (Platelet) |
| TGFBR1 | Transforming Growth Factor, Beta Receptor 1 |
| TIE1 | Tyrosine Kinase With Immunoglobulin-like And Egf-like Domains 1 |
| TIMP3 | TIMP Metallopeptidase Inhibitor 3 |
| TNFRSF1B | Tumor Necrosis Factor Receptor Superfamily, Member 1B |
| TNFSF15 | Tumor Necrosis Factor (Ligand) Superfamily, Member 15 |
| VCAM1 | Vascular Cell Adhesion Molecule 1 |

Table (S3) Genes selected following gene expression analysis to generate the transcriptomic signature of irradiation (continued).

| Protein/ Gene ID | Full name |
|---|---|
| B3GNT2 | UDP-GlcNAc:betaGal beta-1,3-N-acetylglucosaminyltransferase 2 |
| B4GALT5 | UDP-Gal:betaGlcNAc beta 1,4- galactosyltransferase, polypeptide 5 |
| FUCA1 | fucosidase, alpha-L- 1, tissue |
| FUT8 | fucosyltransferase 8 (alpha (1,6) fucosyltransferase) |
| GALNT12 | UDP-N-acetyl-alpha-D-galactosamine:polypeptide N-acetylgalactosaminyltransferase 12 (GalNAc-T12) |
| GALNT14 | UDP-N-acetyl-alpha-D-galactosamine:polypeptide N-acetylgalactosaminyltransferase 14 (GalNAc-T14) |
| GCNT3 | glucosaminyl (N-acetyl) transferase 3, mucin type |
| GCNT4 | glucosaminyl (N-acetyl) transferase 4, core 2 |
| GNPTAB | N-acetylglucosamine-1-phosphate transferase, alpha and beta subunits |
| MAN1C1 | mannosidase, alpha, class 1C, member 1 |
| MAN2A1 | mannosidase, alpha, class 2A, member 1 |
| MGAT5 | mannosyl (alpha-1,6-)-glycoprotein beta-1,6-N-acetyl-glucosaminyltransferase |
| ST8SIA6 | ST8 alpha-N-acetyl-neuraminide alpha-2,8-sialyltransferase 6 |

Table (S4) Genes involved in glycosylation included in the transcriptomic signature of irradiation.

| Protein/ Gene ID | Full name |
|---|---|
| ACTA2 | alpha smooth muscle actin |
| VWF | Von Willebrand Factor |
| CDH5 | VE-Cadherin |
| TAGLN | Smooth muscle 22 alpha |
| TNC | Tenascine C |
| VTN | Vitronectine |
| SNAIL2 | Snail family zing finger 2 (slug) |
| HEY2 | Hairy/enhancer-of-split related with YRPW motif protein 2 |
| SPP1 | Osteopontin |
| TIE1 | tyrosine kinase with immunoglobulin-like and EGF-like domains 1 |
| TEK | tyrosine kinase with immunoglobulin-like and EGF-like domains 2 |
| SERPINE1 | Plasminogene activator inhibitor type 1 |

Table (S5) Genes involved in endothelium-to-mesenchyme transition (EndoMT) included in the transcriptomic signature of irradiation.

| Protein/ Gene ID | Full name |
|---|---|
| CDKN2A | cyclin-dependent kinase inhibitor 2A |
| CDKN1A | cyclin-dependent kinase inhibitor 1A (p21, Cip1) |
| GADD45A | growth arrest and DNA-damage-inducible, alpha |

Table (S6) Genes involved in cellular senescence included in the transcriptomic signature of irradiation.

| Protein/ Gene ID | Full name |
|---|---|
| HIF1A | hypoxia inducible factor 1, alpha subunit (basic helix-loop-helix transcription factor) |
| RELA | v-rel avian reticuloendotheliosis viral oncogene homolog A |
| NFKB1 | nuclear factor of kappa light polypeptide gene enhancer in B-cells 1 |
| SP1 | Sp1 transcription factor |
| CD40 | CD40 molecule, TNF receptor superfamily member 5 |
| CXCL12 | chemokine (C-X-C motif) ligand 12 |
| PDGFA | platelet-derived growth factor alpha polypeptide |
| IL17A | interleukin 17A |
| IFNG | interferon, gamma |
| TGFB1 | transforming growth factor, beta 1 |
| TNF | tumor necrosis factor |

Table (S7) Genes highlighted by sub-network enrichment for common regulators of proteomic and transcritomic data included in the transcriptomic signature of irradiation.

| Protein/ Gene ID | Full name |
|---|---|
| IL2 | interleukin 2 |
| IL4 | interleukin 4 |
| IL10 | interleukin 10 |
| CCL2 | chemokine (C-C motif) ligand 2 |
| CCL4 | chemokine (C-C motif) ligand 4 |

Table (S8) Genes chemokines involved in the response to radiation included in the transcriptomic signature of irradiation.

| Protein/ Gene ID | Full name |
|---|---|
| 18S | |
| ACTB | actin, beta |
| PGK1 | phosphoglycerate kinase 1 |
| MAPK8 | mitogen-activated protein kinase 8 |
| SKI | v-ski avian sarcoma viral oncogene homolog |

Table (S9) Reference genes included in the transcriptomic signature of irradiation.

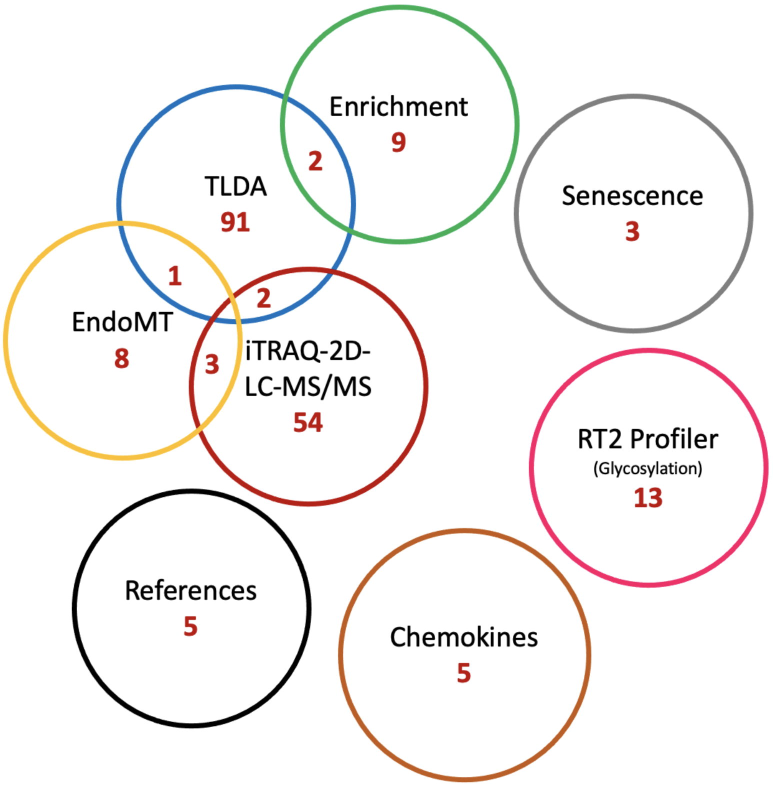


Figure (S1) Summary of the number of genes selected for signature according to the technique or method used, or the cellular process to which they belong. Some genes were highlighted by several means, which is indicated by the overlap between the groups.

| | 1 | 2 | 3 | 4 | 5 | 6 | 7 | 8 | 9 | 10 | 11 |
|---|---|---|---|---|---|---|---|---|---|---|---|
| **1** | 0 | 0 | 1 | 0 | 0 | 0 | 0 | 0 | 0 | 1 | 1 |
| **2** | 0 | 0 | 1 | 0 | 0 | 0 | 0 | 0 | 0 | 0 | 1 |
| **3** | 0 | 0 | 0 | 0 | 1 | 1 | 0 | 0 | 0 | 0 | 0 |
| **4** | 0 | 0 | 1 | 0 | 0 | 0 | 1 | 1 | 1 | 0 | 0 |
| **5** | 0 | 0 | 0 | 0 | 0 | 1 | 0 | 0 | 1 | 0 | 1 |
| **6** | 0 | 0 | 0 | 0 | 0 | 0 | 0 | 0 | 0 | 0 | 1 |
| **7** | 0 | 0 | 0 | 1 | 0 | 0 | 0 | 0 | 1 | 0 | 1 |
| **8** | 0 | 0 | 1 | 0 | 0 | 0 | 0 | 0 | 0 | 1 | 1 |
| **9** | 1 | 0 | 1 | 0 | 1 | 0 | 1 | 1 | 0 | 1 | 0 |
| **10** | 0 | 0 | 0 | 0 | 0 | 0 | 0 | 0 | 1 | 0 | 1 |
| **11** | 0 | 0 | 1 | 0 | 0 | 0 | 1 | 1 | 1 | 0 | 0 |

Table (S10) Example of an adjacency matrix for a directed network.

| | | SARRP clusters | | | | |
|---|---|---|---|---|---|---|
| | | **1** | **2** | **3** | **4** | **5** |
| LINAC clusters | **1** | 13 | 4 | 7 | 2 | 0 |
| | **2** | 1 | 4 | 4 | 2 | 4 |
| | **3** | 11 | 0 | 14 | 2 | 2 |
| | **4** | 3 | 4 | 2 | 11 | 11 |
| | **5** | 0 | 1 | 1 | 8 | 38 |

Table (S11) Contingency table for clusters obtained for LINAC and SARRP.

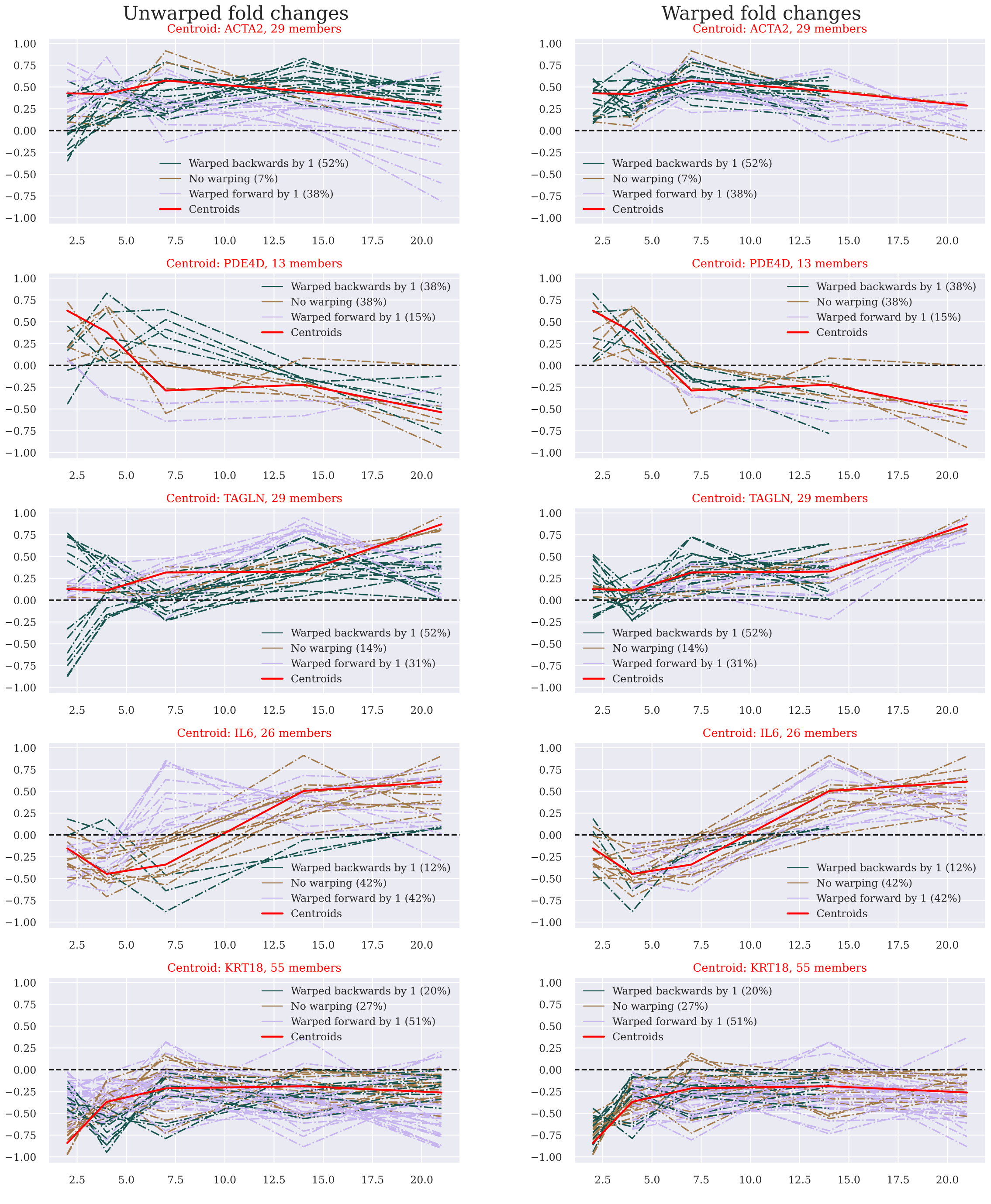


(a) . (b)

Figure (S2) Clustering of the SARRP dataset with $\widehat{\textbf{diss}}$ k-medoids in 5 clusters. a) Means of original normalized fold changes (unaligned). b) Means of warped normalized fold changes (aligned).

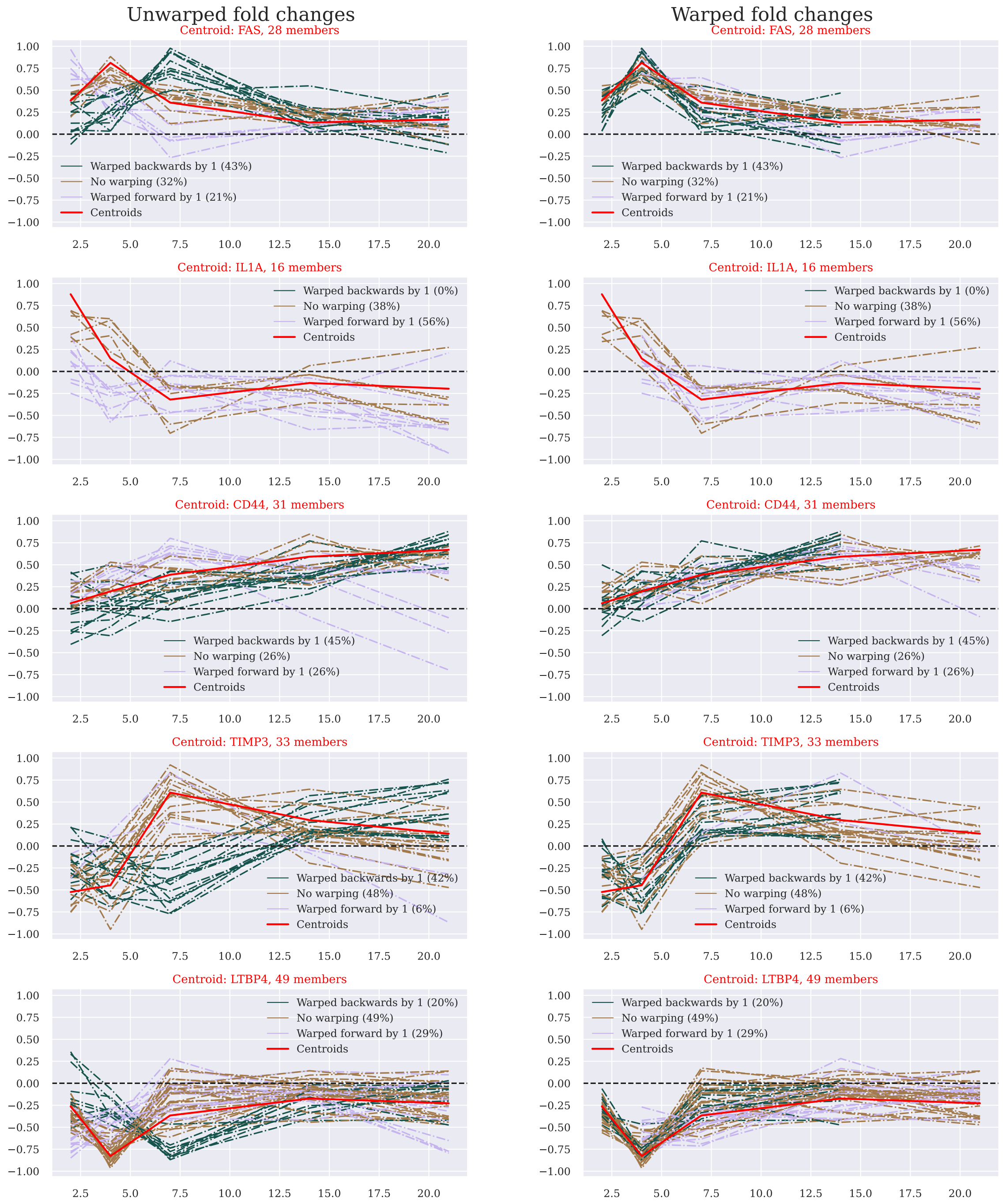


(a) (b)

Figure (S3) Clustering of the LINAC dataset with $\widehat{\textbf{diss}}$ k-medoids in 5 clusters. a) Means of original normalized fold changes (unaligned). b) Means of warped normalized fold changes (aligned).

The following behavior types can be distinguished (top to bottom): up-regulated during the first week and positively tending towards zero, up-regulated during the first week and negatively tending towards zero, steady growth, down-regulated during the first week and positively tending towards zero, down-regulated during the first week and negatively tending towards zero.

| Sparsity | 1-2 | 1-3 | 1-4 | 1-5 | 2-3 | 2-4 | 2-5 | 3-4 | 3-5 | 4-5 |
|---|---|---|---|---|---|---|---|---|---|---|
| 0.6 | 0.0409 | 0.0043 | 0.0338 | 0.1641 | 0.0164 | 0.0924 | 0 | 0.0106 | 0.9532 | 0.0001 |
| 0.625 | 0.0177 | 0.0262 | 0.0301 | 0.2275 | 0.0254 | 0.0588 | 0 | 0.0072 | 0.6112 | 0 |
| 0.65 | 0.0174 | 0.0107 | 0.0055 | 1 | 0.0085 | 0.2453 | 0 | 0.0265 | 1 | 0 |
| 0.675 | 0.037 | 0.0234 | 0.0016 | 1 | 0.0047 | 0.1811 | 0 | 0.1123 | 0.91 | 0 |
| 0.7 | 0.0167 | 0.1524 | 0.0001 | 1 | 0.008 | 0.4579 | 0.0001 | 0.0614 | 0.6945 | 0 |
| 0.725 | 0.0239 | 0.2983 | 0.0001 | 1 | 0.0094 | 0.0965 | 0.001 | 0.0242 | 0.4636 | 0 |
| 0.75 | 0.0415 | 0.4741 | 0.0009 | 1 | 0.0074 | 0.2289 | 0.0101 | 0.0711 | 1 | 0 |
| 0.775 | 0.0219 | 0.6811 | 0.0008 | 1 | 0.033 | 0.1536 | 0.1077 | 0.1422 | 0.3457 | 0.0001 |
| 0.8 | 0.005 | 0.5479 | 0.0006 | 1 | 0.0339 | 0.0682 | 0.1304 | 0.3585 | 0.2193 | 0.0001 |
| 0.825 | 0.0294 | 0.8631 | 0.0007 | 1 | 0.1298 | 0.1281 | 0.1557 | 0.2247 | 0.9624 | 0.0005 |
| 0.85 | 0.0656 | 0.818 | 0.0007 | 1 | 0.6126 | 0.3664 | 0.4383 | 0.1429 | 1 | 0.0043 |
| 0.875 | 0.0962 | 0.7424 | 0.002 | 1 | 1 | 0.6016 | 0.6775 | 0.5252 | 1 | 0.0406 |
| 0.9 | 0.1236 | 0.6786 | 0.0165 | 1 | 0.3868 | 0.9705 | 0.2095 | 0.1479 | 0.615 | 0.0571 |

Table (S12) P-values associated with the results of the Barnard exact test performed on every connection (rows) of mesoscopic graphs obtained for LINAC and SARRP constructed with a given sparsity level (columns).

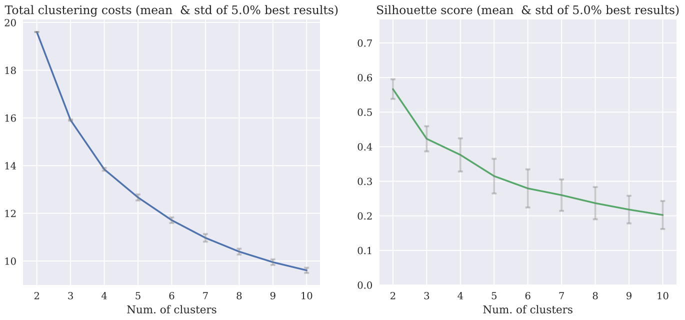


Figure (S4) Means of cost and silhouette score with standard deviation of the 20% best outcomes of clustering of the LINAC dataset with $\widehat{\mathbf{diss}}$ k-medoids for the numbers of clusters in the set of fold changes ranging from 2 to 10. The most distinguishable shoulder for the total cost can be observed for 3 clusters, whereas the silhouette score declines for larger number of clusters. Both criteria suggest that smallest number of clusters should be chosen.

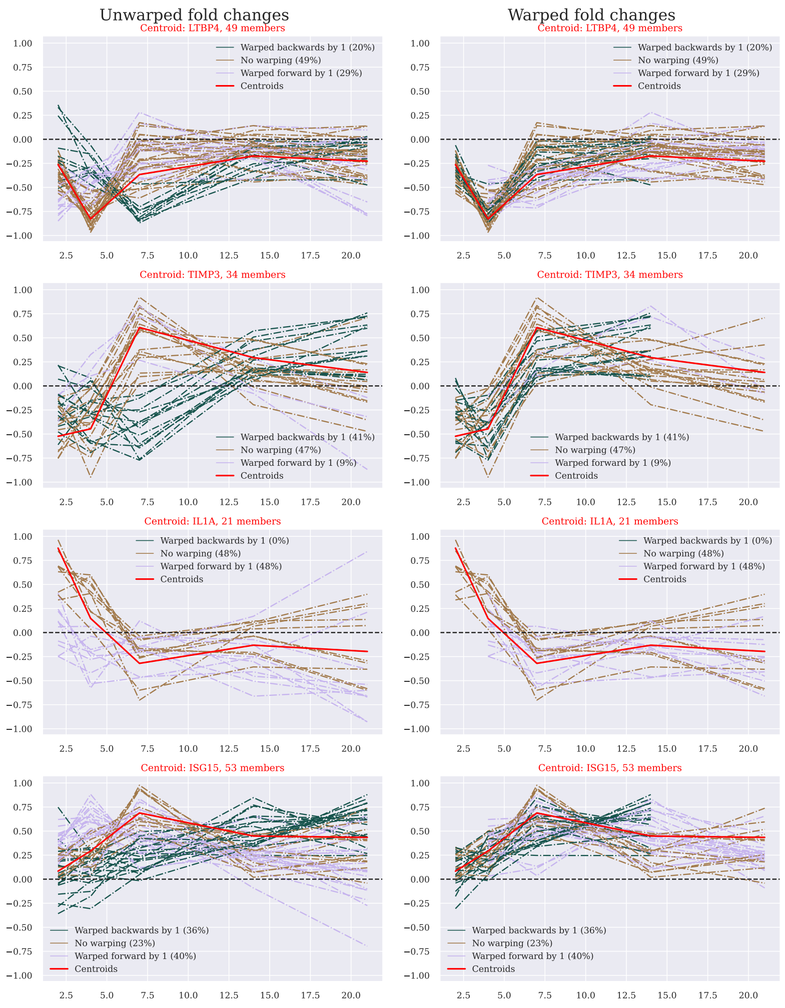


(a) (b)

Figure (S5) Clustering of the LINAC dataset with $\widehat{\mathbf{diss}}$ k-medoids in 4 clusters. a) Means of original normalized fold changes (unaligned). b) Means of warped normalized fold changes (aligned).

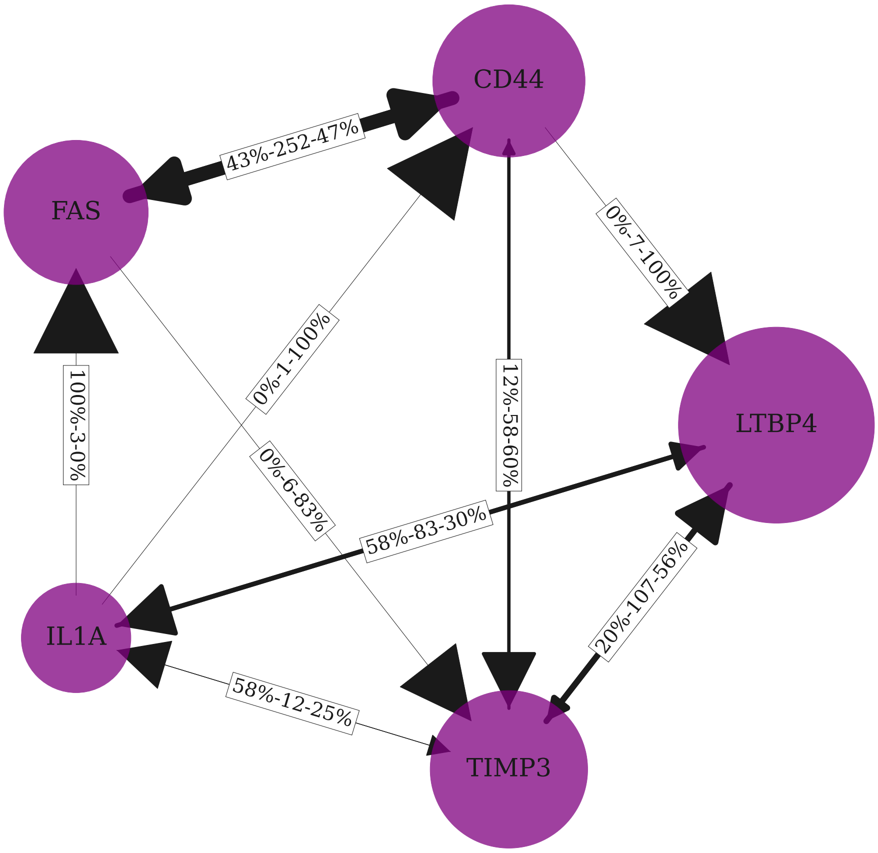


Figure (S6) Mesoscopic view of the LINAC-specific fold changes network. Nodes represent clusters, labeled with their centroid genes, with node sizes corresponding to respective cluster sizes, and edges summarizing the connections between clusters based on the original adjacency matrix. Edge thickness corresponds to the total number of connections, arrow size to the proportion of predictive connections of the corresponding type. Edge labels read as follows: *"% of connections right → left - Total number of connections - % of connections left → right"*.

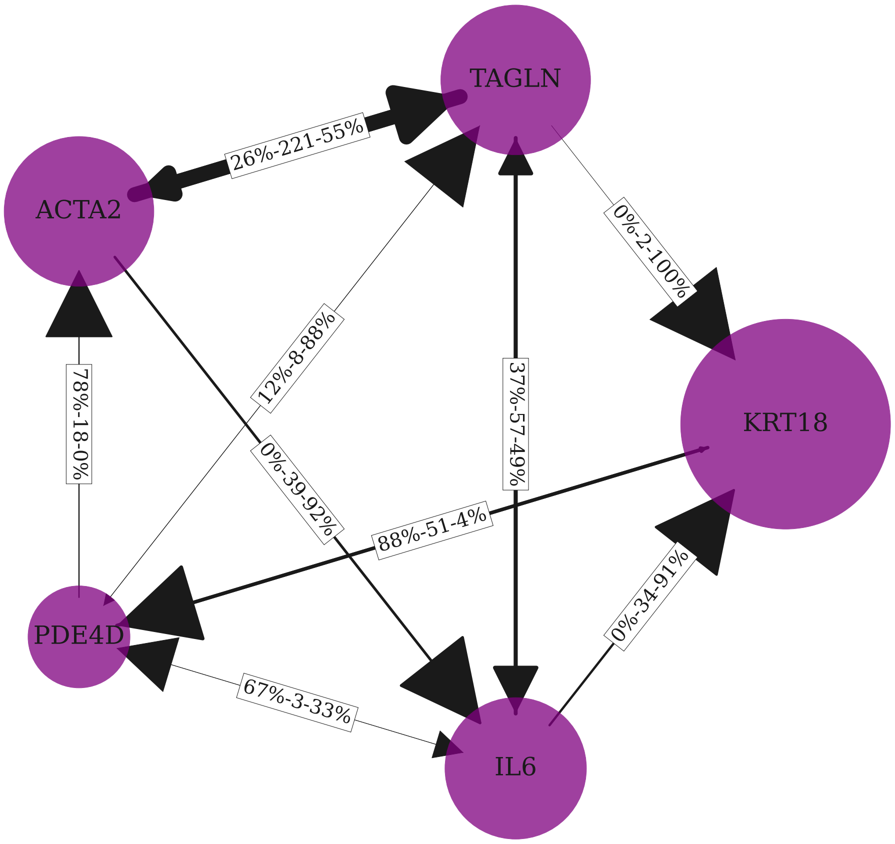


Figure (S7) Mesoscopic view of the SARRP-specific fold changes network. Nodes represent clusters, labeled with their centroid genes, with node sizes corresponding to respective cluster sizes, and edges summarizing the connections between clusters based on the original adjacency matrix. Edge thickness corresponds to the total number of connections, arrow size to the proportion of predictive connections of the corresponding type. Edge labels read as follows: *"% of connections right → left - Total number of connections - % of connections left → right"*.

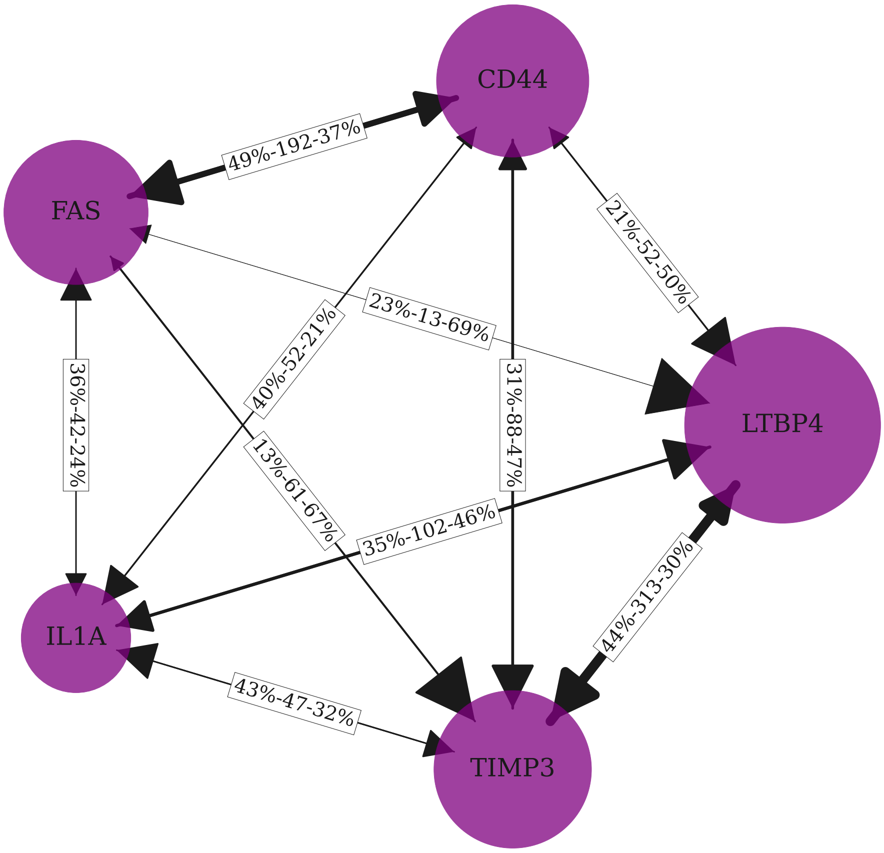


Figure (S8) Mesoscopic view of the hybrid fold changes network. Nodes represent clusters (LINAC), labeled with their centroid genes, with node sizes corresponding to respective cluster sizes, and edges summarizing the connections between clusters based on the original adjacency matrix (SARRP). Edge thickness corresponds to the total number of connections, arrow size to the proportion of predictive connections of the corresponding type. Edge labels read as follows: *"% of connections right → left - Total number of connections - % of connections left → right"*.

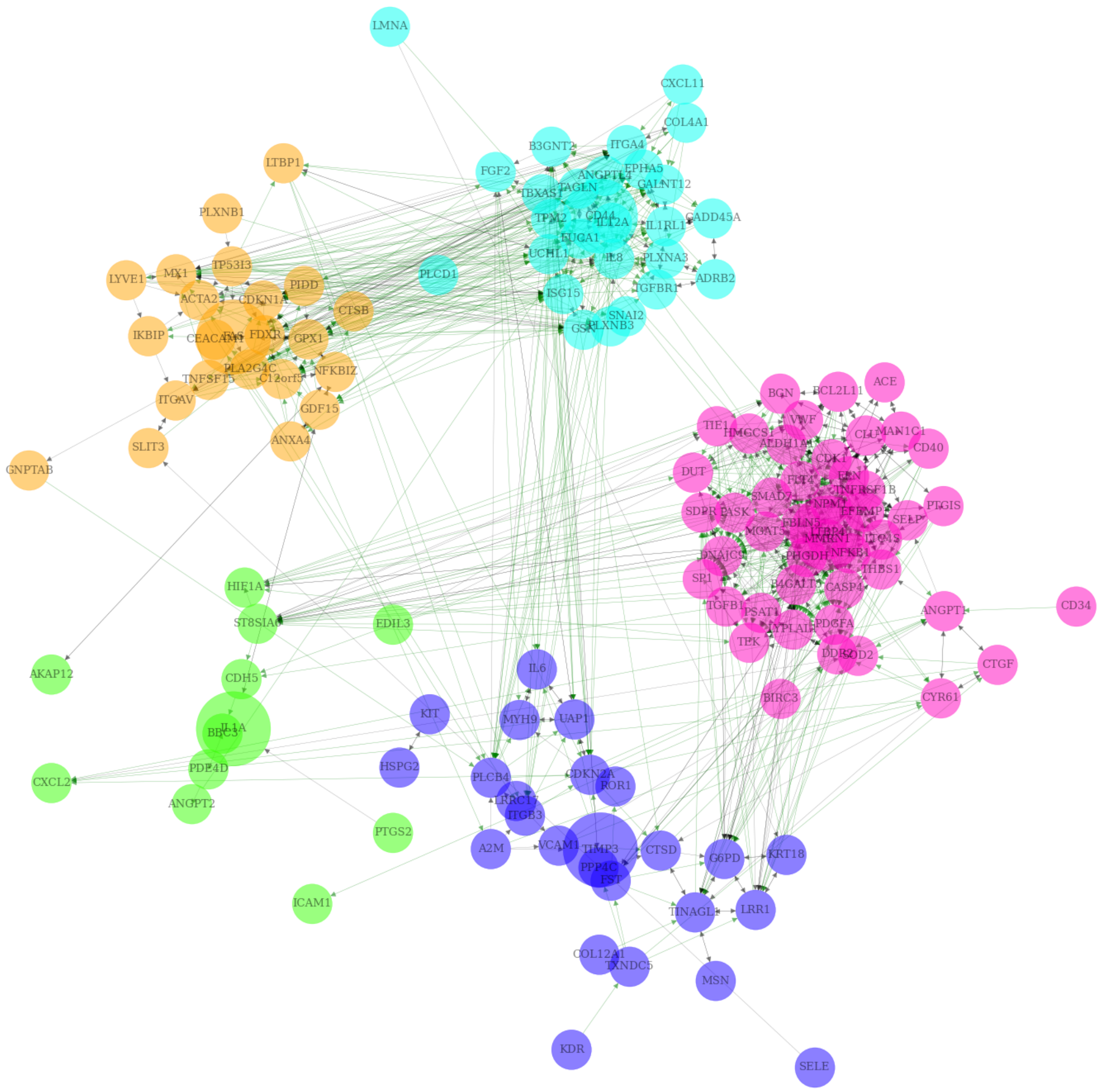


Figure (S9) Microscopic view of the LINAC directed fold changes network. Green edges correspond to predictive connections, whereas gray to simultaneous ones. Here the connections are those arising from the intersection of LINAC and SARRP networks, i.e. those that are common for both conditions. The network has a block structure, where blocks denoted by different colors correspond to clusters inferred from the LINAC dataset. Within each block, the fold changes are placed around their centroid (bigger node) according to the Kamada-Kawai method.

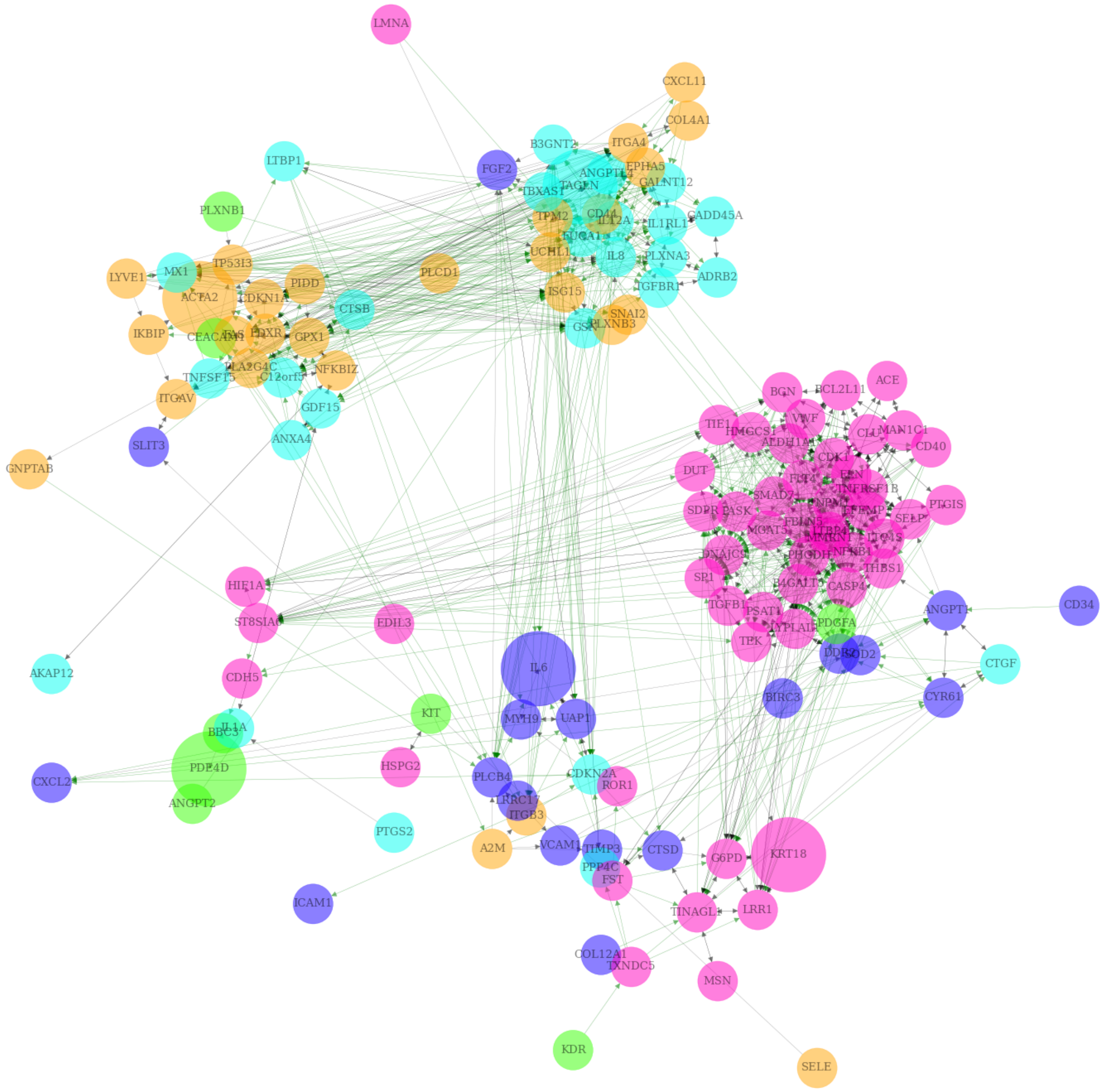


Figure (S10) Microscopic view of the hybrid fold changes network. Green edges correspond to predictive connections, whereas gray to simultaneous ones. Here the connections are those arising from the intersection of LINAC and SARRP networks, i.e. those that are common for both conditions. The network has a (hybrid) block structure. The blocks in terms of layout correspond to clusters inferred from the LINAC dataset, the node positions are the same as those in the LINAC network view (Figure S9). The cluster colors match those of the LINAC network but are assigned according to the SARRP clustering. The bigger nodes are the centroids with respect to the SARRP clustering.

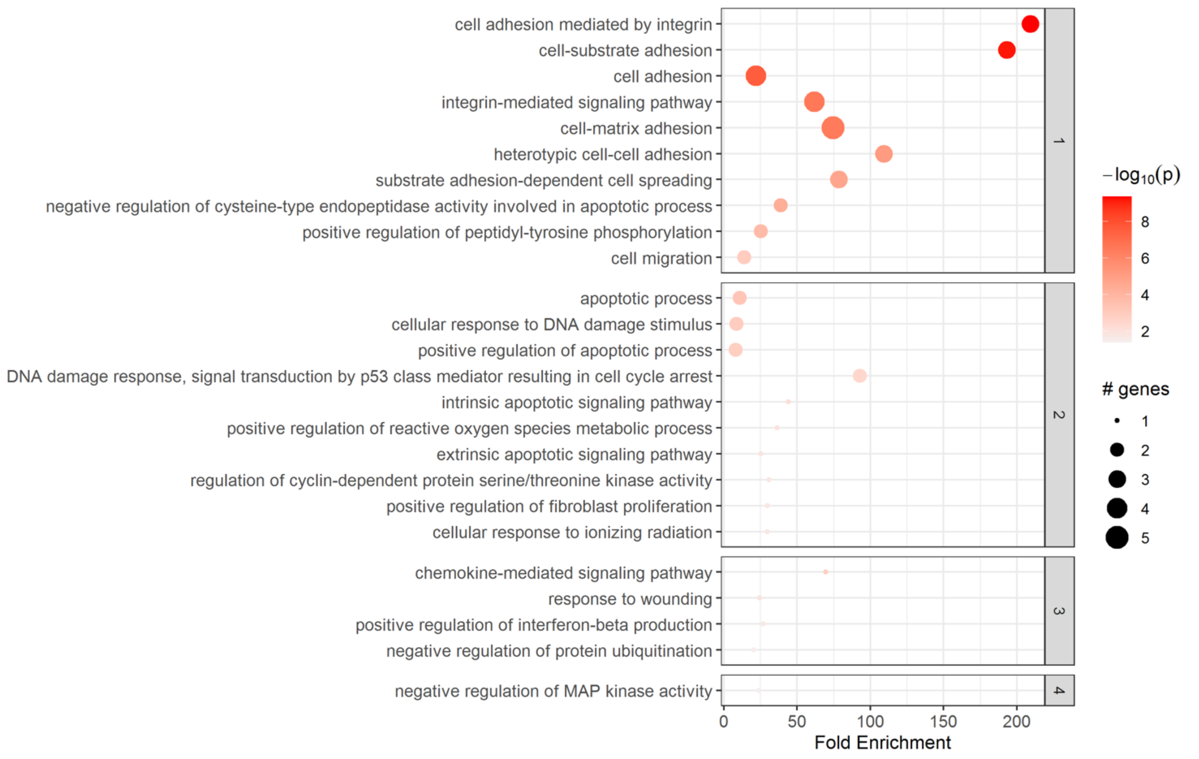


Figure (S11) Results of the enrichment analysis performed with Pathfinder on the genes from cluster 1 of the SARRP dataset.

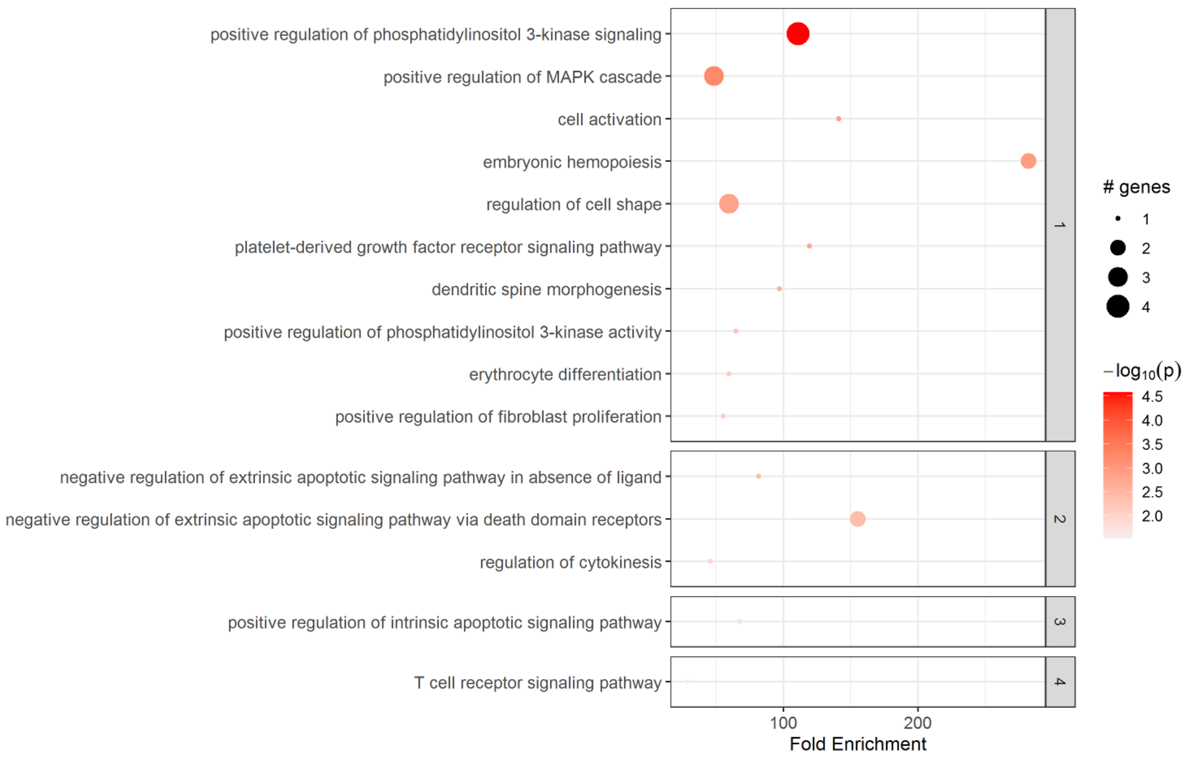


Figure (S12) Results of the enrichment analysis performed with Pathfinder on the genes from cluster 2 of the SARRP dataset.

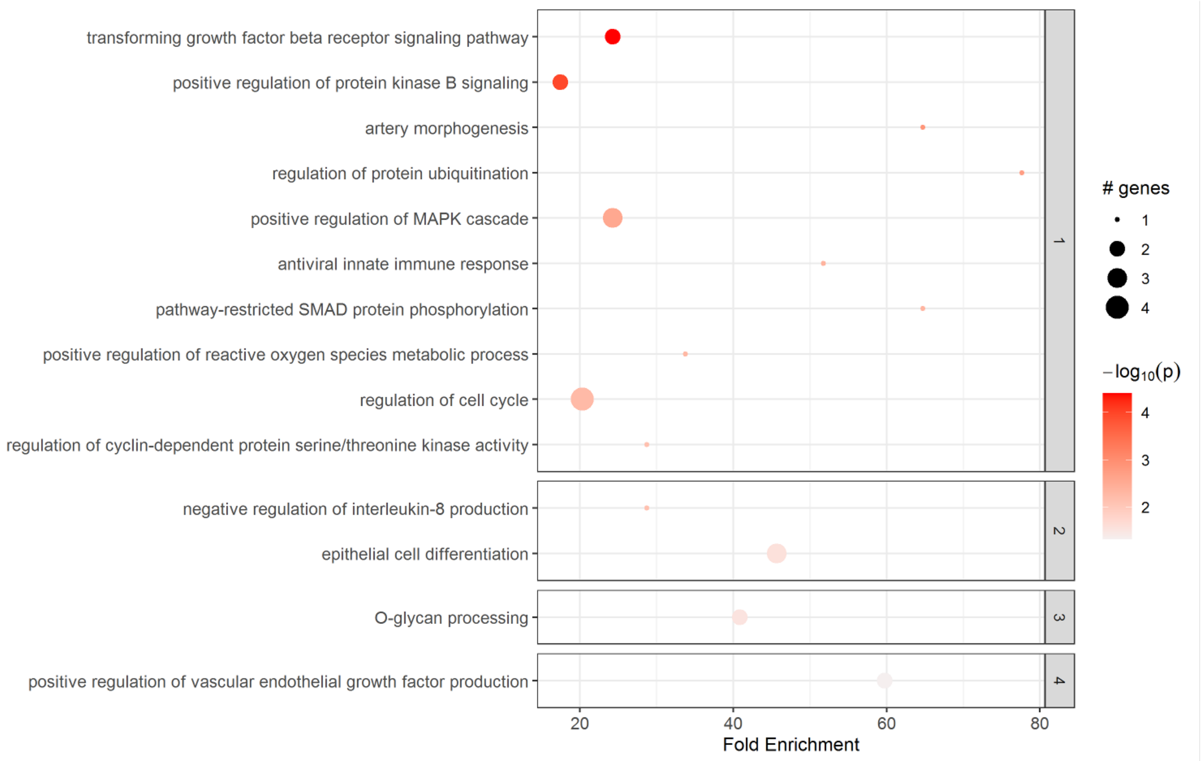


Figure (S13) Results of the enrichment analysis performed with Pathfinder on the genes from cluster 3 of the SARRP dataset.

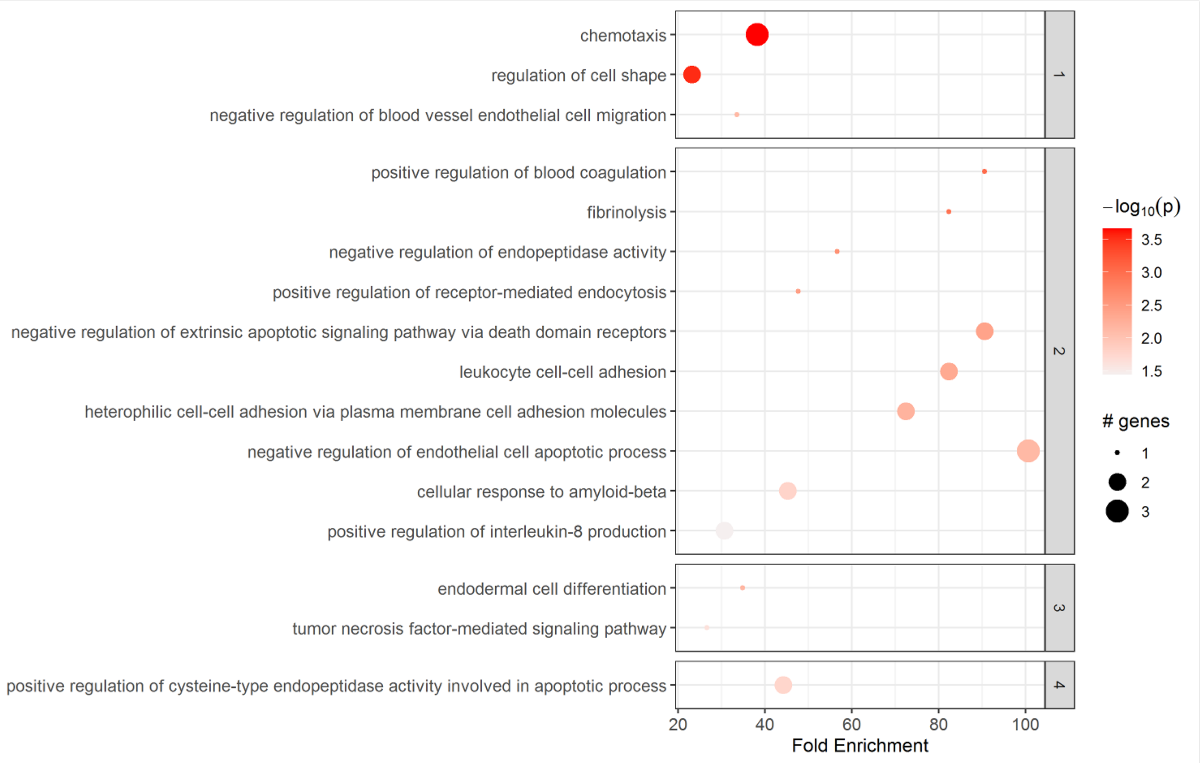


Figure (S14) Results of the enrichment analysis performed with Pathfinder on the genes from cluster 4 of the SARRP dataset.

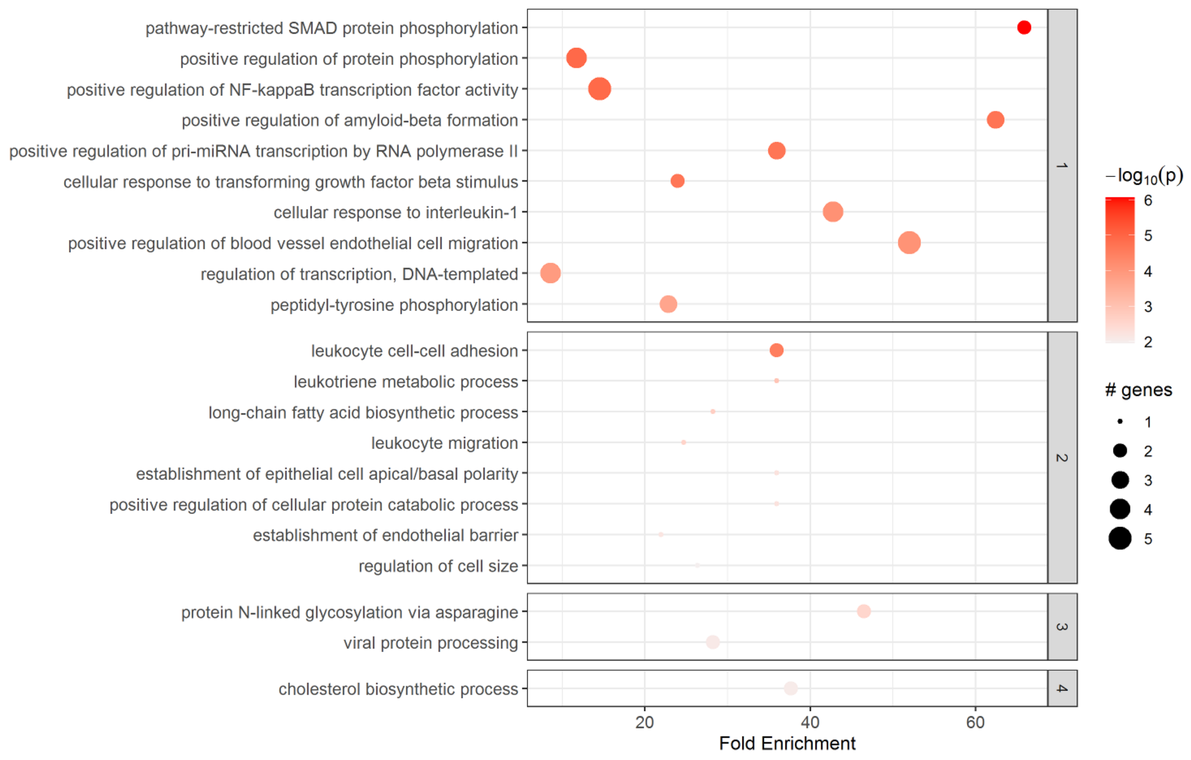


Figure (S15) Results of the enrichment analysis performed with Pathfinder on the genes from cluster 5 of the SARRP dataset.

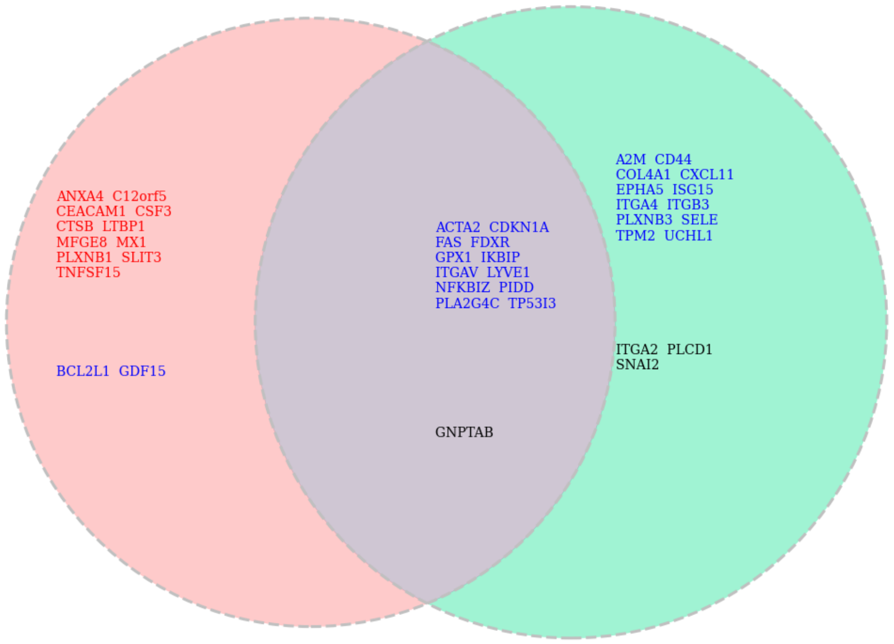


Figure (S16) Venn diagrams illustrating the distribution of genes of cluster 1 between LINAC (left) and SARRP (right). Red indicates genes that have been assigned to this cluster by all methods (k-medoids, UMAP and SBM), blue indicates those that have been assigned to this cluster by two methods out of three, and black the remaining genes.

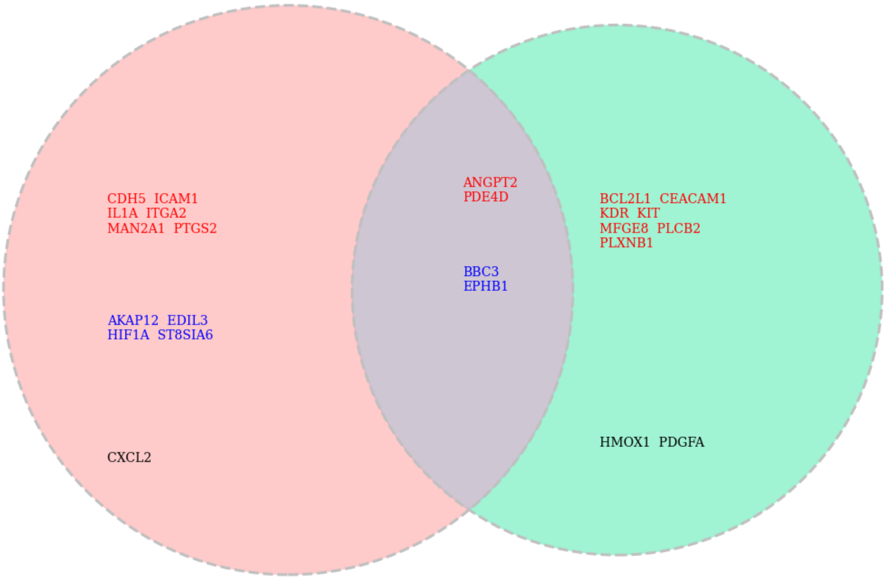


Figure (S17) Venn diagrams illustrating the distribution of genes of cluster 2 between LINAC (left) and SARRP (right). Red indicates genes that have been assigned to this cluster by all methods (k-medoids, UMAP and SBM), blue indicates those that have been assigned to this cluster by two methods out of three, and black the remaining genes.

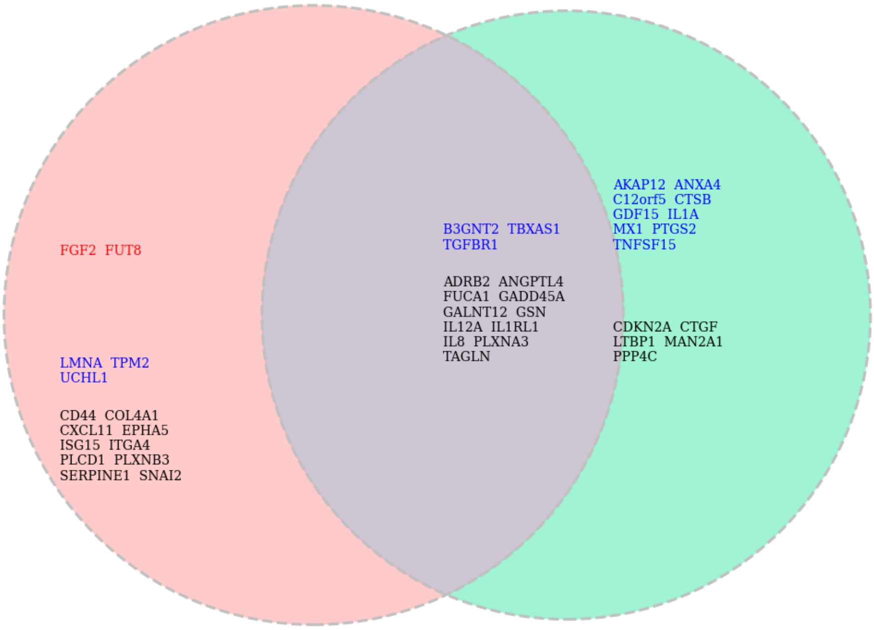


Figure (S18) Venn diagrams illustrating the distribution of genes of cluster 3 between LINAC (left) and SARRP (right). Red indicates genes that have been assigned to this cluster by all methods (k-medoids, UMAP and SBM), blue indicates those that have been assigned to this cluster by two methods out of three, and black the remaining genes.

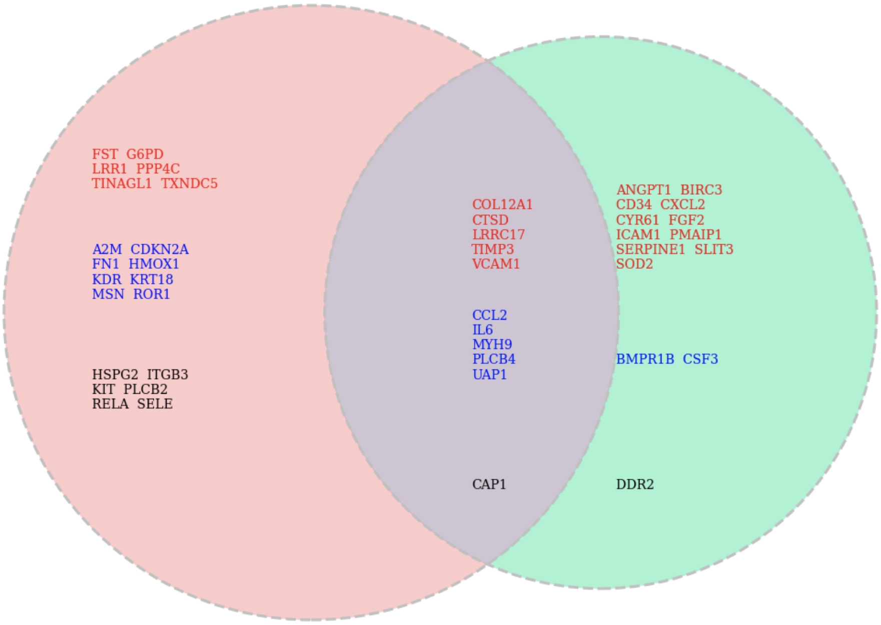


Figure (S19) Venn diagrams illustrating the distribution of genes of cluster 4 between LINAC (left) and SARRP (right). Red indicates genes that have been assigned to this cluster by all methods (k-medoids, UMAP and SBM), blue indicates those that have been assigned to this cluster by two methods out of three, and black the remaining genes.

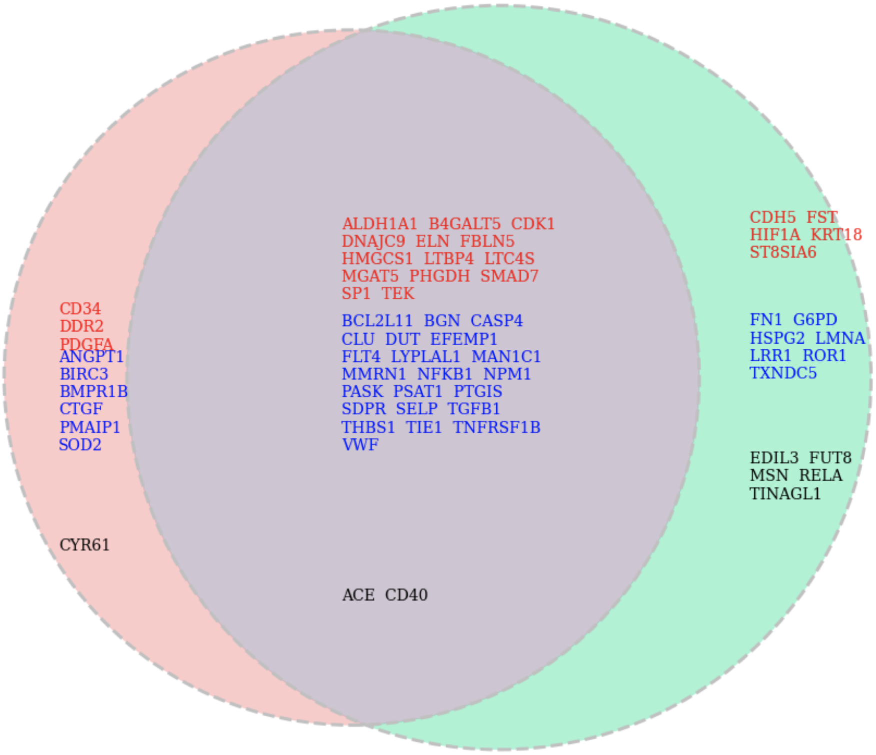


Figure (S20) Venn diagrams illustrating the distribution of genes of cluster 5 between LINAC (left) and SARRP (right). Red indicates genes that have been assigned to this cluster by all methods (k-medoids, UMAP and SBM), blue indicates those that have been assigned to this cluster by two methods out of three, and black the remaining genes.

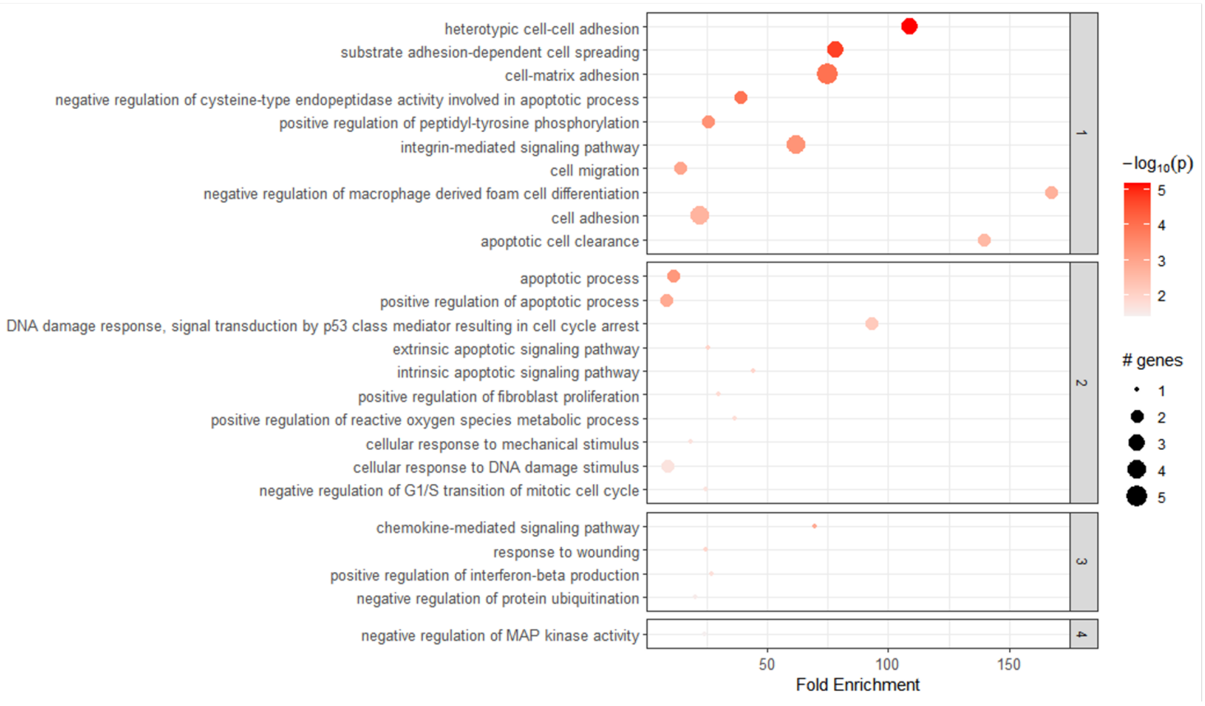


Figure (S21) Results of the enrichment analysis performed with Pathfinder on the genes from cluster 1 of the LINAC dataset.

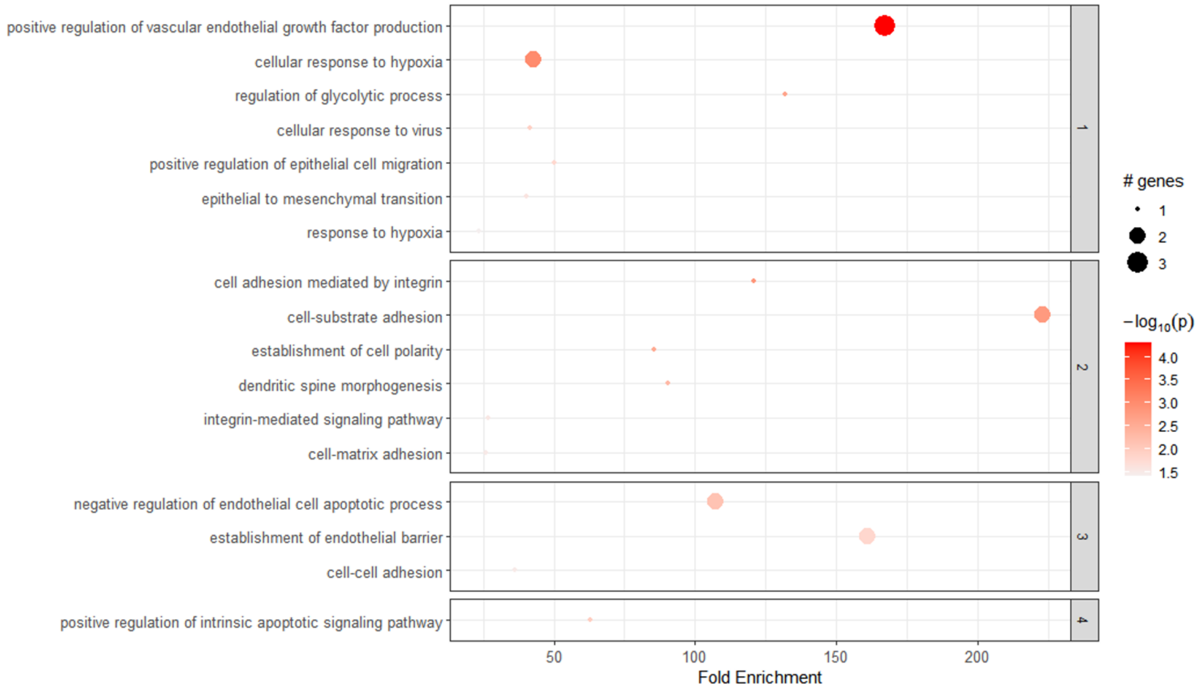


Figure (S22) Results of the enrichment analysis performed with Pathfinder on the genes from cluster 2 of the LINAC dataset.

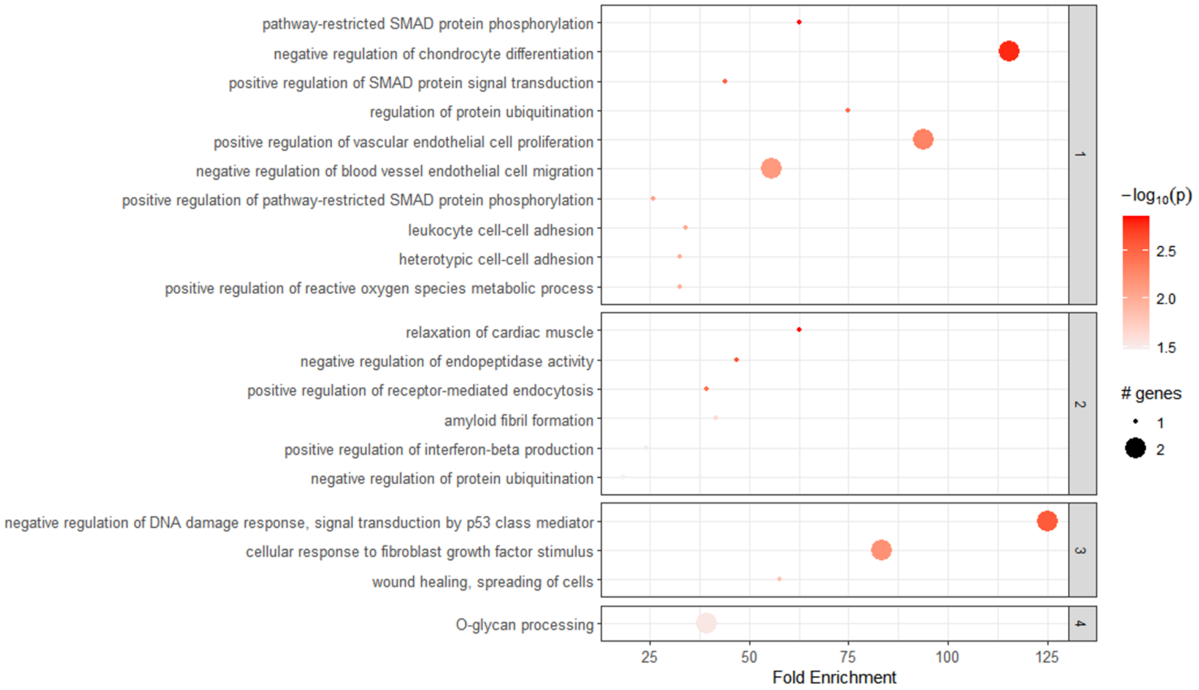


Figure (S23) Results of the enrichment analysis performed with Pathfinder on the genes from cluster 3 of the LINAC dataset.

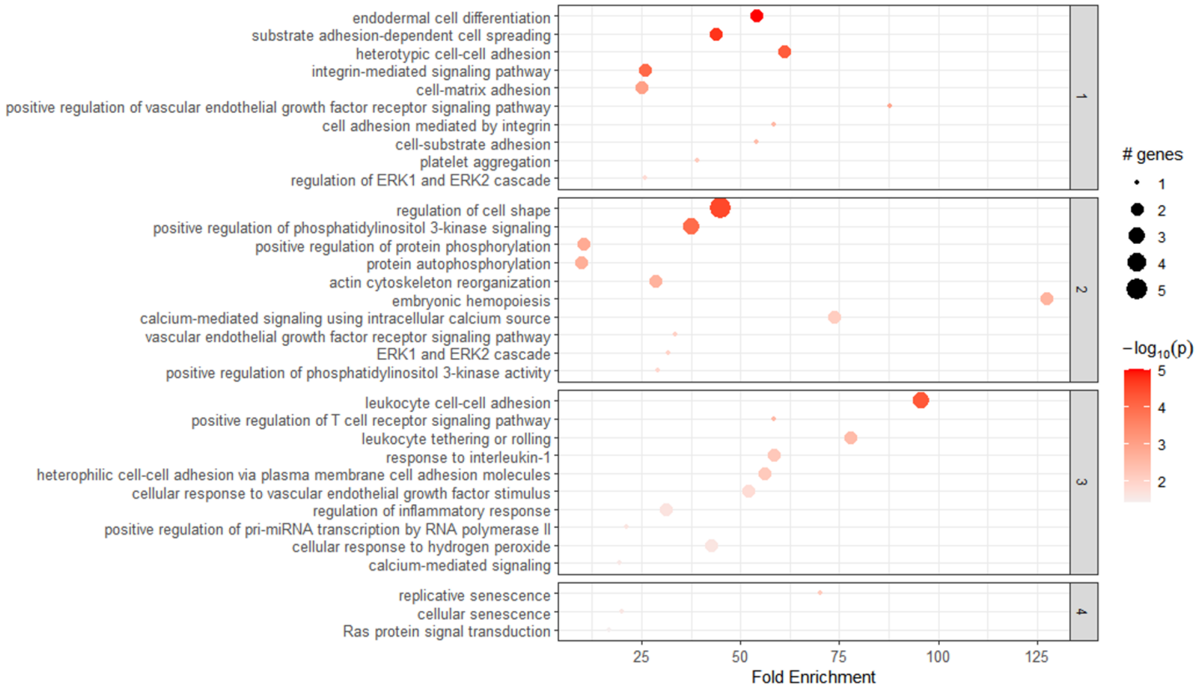


Figure (S24) Results of the enrichment analysis performed with Pathfinder on the genes from cluster 4 of the LINAC dataset.

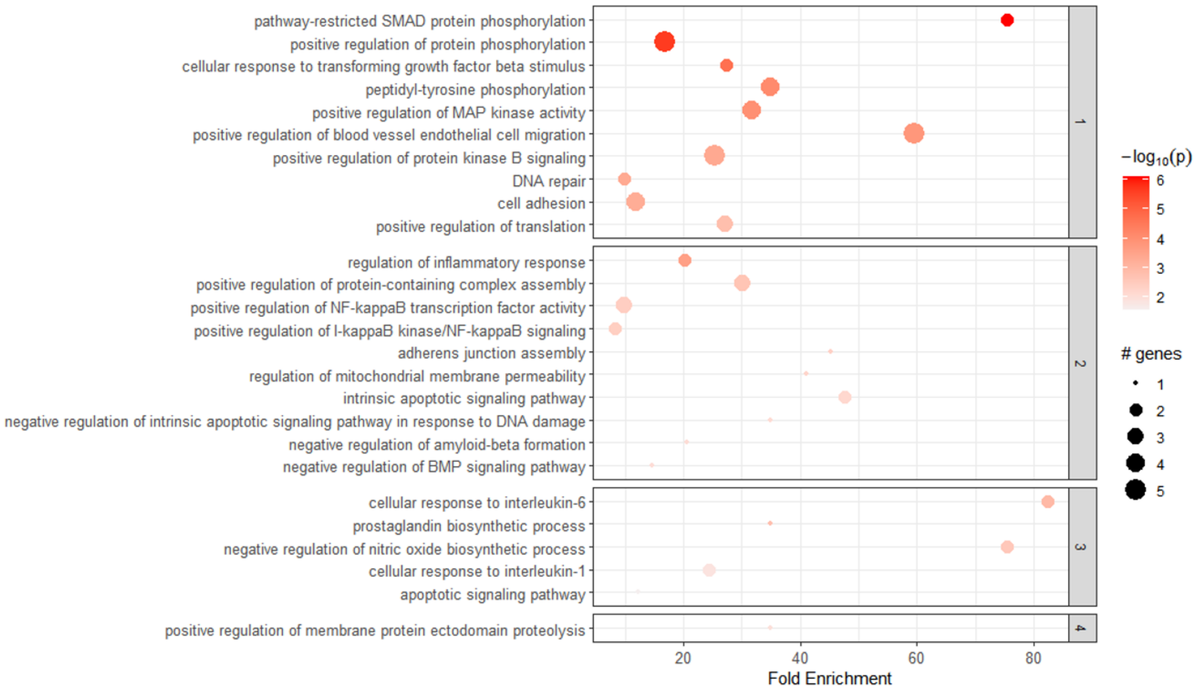


Figure (S25)

Results of the enrichment analysis performed with Pathfinder on the genes from cluster 5 of the LINAC dataset.
